# Invasive cancer cells soften collagen networks and disrupt stress-stiffening via volume exclusion, contractility and adhesion

**DOI:** 10.1101/2025.04.11.648338

**Authors:** Irène Nagle, Margherita Tavasso, Ankur D. Bordoloi, Iain A. A. Muntz, Gijsje H. Koenderink, Pouyan E. Boukany

**Author notes:** Corresponding authors: Gijsje H. Koenderink, Pouyan E. Boukany. These authors contributed equally to this work (Irène Nagle, Margherita Tavasso).

## Abstract

Collagen networks form the structural backbone of the extracellular matrix in both healthy and cancerous tissues, exhibiting nonlinear mechanical properties that crucially regulate tissue mechanics and cell behavior. Here, we investigate how the presence of invasive breast cancer cells (MDA-MB-231) influences the polymerization kinetics and mechanics of collagen networks using bulk shear rheology and rheo-confocal microscopy. We show that embedded cancer cells delay the onset of collagen polymerization due to volume exclusion effects. During polymerization, the cells (at 4% volume fraction) cause an unexpected time-dependent softening of the network. We show that this softening effect arises from active remodeling via adhesion and contractility rather than from proteolytic degradation. At higher cell volume fractions, the dominant effect of the cells shifts to volume exclusion, causing a two-fold reduction of network stiffness. Additionally, we demonstrate that cancer cells suppress the characteristic stress-stiffening response of collagen. This effect (partially) disappears when cell adhesion and contractility are inhibited, and it is absent when the cells are replaced by passive hydrogel particles. These findings provide new insights into how active inclusions modify the mechanics of fibrous networks, contributing to a better understanding of the role of cells in the mechanics of healthy and diseased tissues like invasive tumors.

## 1. Introduction

Biological tissues exhibit tissue-specific mechanical properties, which arise from the complex interplay between extracellular matrix (ECM) composition, cell activity, and their relative proportions [1]. The ECM is a fibrous network composed of different biopolymers, including collagen, fibrin, and fibronectin. It serves as a scaffold for cell adhesion and controls the shape and mechanical properties of the tissue [2]. Additionally, it provides biochemical and biophysical cues to cells, which regulate essential processes such as cellular differentiation, tissue morphogenesis, wound healing, and homeostasis [3, 4]. Conversely, the ECM can also be altered by the activity of the cells themselves, including active force application [5] as well as chemical modifications, such as proteolytic degradation of the matrix [6]. Mechanochemical feedback mechanisms ensure tissue homeostasis and overall health [7] and disruptions in this feedback can contribute to pathological conditions such as fibrosis [8] or cancer [9]. These pathological states involve matrix remodeling and alterations of the tissue mechanics, at both local and global scales. In the context of cancer, for example, the tumor microenvironment undergoes extensive remodeling due to the activity of various cell types, including cancer-associated fibroblasts and malignant cancer cells. This remodeling generally results in increased tissue stiffness, primarily driven by ECM deposition, crosslinking, and contractile forces exerted by cells [9, 10].

A major challenge in deciphering the contributions of cells and ECM to the overall tissue mechanics arises from the intrinsic complexity of biological tissues, which feature heterogeneous compositions and architectures. One successful approach to address this limitation has been the development of biomimetic tissue models, where ECM components, such as collagen networks, are populated with cells, in order to replicate natural tissues [11, 12]. Collagen is the main determinant of the overall mechanical properties of connective tissues [13] and exhibits highly nonlinear responses to shear or tensile loading, with stiffness increasing sharply as strain increases [14]. Cells within the fibrous ECM network can act as volume-conserving inclusions, modifying its mechanical properties in a similar way to inert particles [15]. Recently, there have been several studies on the effect of inert microparticles that serve as passive inclusions on the mechanical properties of fibrous networks [15, 16, 17]. Stiff inclusions at high volume fractions can lead to compression-stiffening due to stretching of the interstitial ECM network. Nevertheless, cells are different from passive inclusions, as they actively interact with the surrounding network. Among cellular inclusions, fibroblasts have been extensively studied due to their ability to generate prestress and bundle ECM fibers. Their contractile forces stiffen the ECM by increasing fiber alignment and tension, reinforcing the overall stiffness of the network [18, 19]. For some other ECM components (like fibronectin and fibrin), tension can induce domain unfolding at submicron scales, undetectable by confocal microscopy [20, 21, 22]. However, this is not known to occur for fibrillar collagen. The mechanical influence of other cell types, such as invasive cancer cells, that interact dynamically with the matrix, remains less understood. Cancer cells, which remodel the ECM through matrix proteolysis and matrix deposition, adhesion, and dynamic forces [23, 24], may influence tissue mechanics in a distinct manner. To better understand the role of collagen in regulating the mechanics of cell-ECM composite systems, a two-phase model has been recently proposed, but only in the limit of high cell density cancer aggregates [25]. Despite the critical function of cells in the tumor microenvironment, the precise role of cancer cells in modifying ECM mechanics, particularly in comparison to passive inclusions and fibroblasts, remains largely unexplored. Moreover, a clear disentangling between the contributions of passive mechanical effects versus active cellular processes to tissue mechanics is still lacking.

Here, we address this research gap by investigating how local cell-ECM interactions regulate the global mechanics of collagen networks in a biomimetic tissue model composed of fibrillar collagen type I with embedded cancer cells (volume fractions ranging from 0.4% to 20%). We initially test human dermal fibroblasts and compare their behavior to that of highly invasive MDA-MB-231 breast cancer cells, which are known to remodel and reorganize collagen type I fibers [26]. We explore by bulk shear rheology how the presence of the cells influences collagen polymerization and the mechanical properties of the final network. We show that the cancer cells delay the kinetics of collagen network assembly and cause softening of the collagen network. In addition, they suppress the stress-stiffening response of collagen networks above a threshold volume fraction. These findings are in marked contrast with the classical stiffening response seen with highly contractile fibroblasts. By comparing the impact of cells with that of passive cell-sized microparticles, we show that cells influence collagen polymerization and mechanics through a combination of volume exclusion and active remodeling dependent on integrin-mediated cell-matrix adhesion. Using a custom-built rheo-confocal microscope, we demonstrate that the cells dynamically remodel the collagen network in their local vicinity as it polymerizes. Our results show that cells influence collagen network mechanics through a delicate combination of volume exclusion, cell adhesion and contractility, providing new insights into the physical mechanisms that determine the mechanical properties of healthy and diseased tissues.

## 2. Materials and methods

### 2.1. Cell culture

Human triple negative breast cancer cells (MDA-MB-231) stably transfected with either LifeAct-GFP (used in most experiments) or nls-mCherry (used only in collagen DQ assays to test collagen degradation) were cultured in DMEM high glucose medium (Gibco), supplemented with 10% fetal bovine serum (FBS, Gibco) and 1% antibiotic-antimycotic solution (15240062, Gibco). This serum-enriched medium is referred to as complete medium. Human dermal fibroblasts (HDF, NHDF-Ad, CC-2511, Lonza) were cultured in IMDM medium (31980022, Gibco) supplemented with 10% FBS (Gibco) and 1% Penicillin-Streptomycin (P/S, 15140122, Gibco). All cells were maintained in a 37°C incubator with 5% CO_2_. MDA-MB-231 cells were subcultured at 90-100% confluency, and HDF cells were subcultured at 80% confluency. Cell counts and viability were assessed with the TC20^TM^ Automated Cell Counter (Bio-Rad) and Trypan Blue 0.4% (15250061, Gibco). MDA-MB-231 cells were cultured until passage 15 and human dermal fibroblasts were cultured until passage 11. The absence of mycoplasma was tested at least once every 4 months.

### 2.2. Fabrication of soft hydrogel microparticles

Soft hydrogel microparticles with an average radius of 13 ± 6 *µ*m were manufactured from acrylamide-co-acrylic-acid (PAA) by Rick Rodrigues (T. Schmidt lab, Leiden University) following a procedure described in [27]. The microparticles had an average Young’s modulus of 4.9 ± 1.4 kPa as determined by shallow indentation measurements with an atomic force microscope [27]. They were stored at 4°C in PBS 1x with 1% (volume percentage) sodium azide at a particle volume fraction of ∼ 15%.

### 2.3. Pharmacological inhibition of cell adhesion, contractility and metalloproteinases activity

Non-muscle myosin II activity was inhibited by treating cells with (±)-blebbistatin (ab120425, Abcam), which inhibits the ATPase activity of myosin by blocking entry into the strong binding state [28]. The blebbistatin powder was dissolved in dimethylsulfoxide (DMSO) at a stock concentration of 5 mM. Cells adhering in plastic culture flasks were incubated with 10 *µ*M blebbistatin [29] in complete medium (DMEM, 10% FBS and 1% antibiotic-antimycotic) in the incubator (37°C, 5 %CO_2_) for 3 hours. Cells were then trypsinized (Trypsin-EDTA 0.25%, phenol red, Gibco), centrifuged for 4 minutes at 200 g and resuspended in the desired volume of complete medium, to obtain a final concentration of 10 *µ*M blebbistatin, right before the rheology experiments.

Integrin *β*1-mediated cell adhesion to collagen was inhibited by treating cells with the anti-*β*1 integrin antibody (CD29, clone p5d2, MAB17781, R&D Systems). The antibody, supplied as a lyophilized powder, was dissolved in sterile PBS 1x at a stock concentration of 0.5 mg/mL. After cell detachment by trypsinization, cells were incubated in suspension with 10 *µ*g/mL anti-*β*_1_ integrin antibody for 15 minutes at 37°C. The cells were then centrifuged for 2 minutes at 200 g and the cell pellet was resuspended in the desired volume of complete medium, right before the rheology experiments.

Batimastat (BB-94), a broad spectrum MMP-inhibitor (ab146619, Abcam), was dissolved in DMSO to a stock concentration of 1 mM. Cells adhering in plastic culture flasks were incubated with 10 *µ*M batimastat [30] in complete medium in the incubator for 3 hours. Cells were then trypsinized, centrifuged for 4 minutes at 200 g and resuspended in the desired volume of complete medium, to obtain a final concentration of 10 *µ*M batimastat, right before the rheology experiments.

### 2.4. Preparation of cell– and microparticle-embedded collagen networks

Throughout this work, we used type I atelocollagen extracted from bovine hides, supplied as a 10 mg/mL solution in 0.01 N HCl (FibriCol^®^, 5133, Advanced Biomatrix) and stored at 4°C. The samples were prepared in Eppen-dorf tubes on ice just before experiments, to prevent premature collagen polymerization. First, the pH was adjusted to 7.4 (checked with pH paper strips, Whatman^®^, Cytiva) with 1% (volume percentage) 0.1 M NaOH (Sigma-Aldrich) and the salt concentration was adjusted with PBS 10x (524650-1EA, Merck-Millipore). Next, complete medium was added to reach a final collagen concentration of 4 mg/mL. If applicable, hydrogel microparticles or cells were included in the complete medium, in varying volume fractions. The cellular volumes were calculated from the average cell size measured by flow cytometry (see Figure S1). The cellular volume fractions were determined from the average cellular volume and the cell number density. Each hydrogel formulation was carefully adjusted to maintain constant collagen and salt concentrations in the fluid phase by taking into account the volume occupied by the cells or hydrogel microparticles.

### 2.5. Bulk rheology measurements

Rheology tests were performed with two stress-controlled rotational rheometers (Physica MCR 501, Anton Paar, Graz, Austria) equipped with a parallel plate geometry with a diameter of 20 mm and using a gap of 160 *µ*m. For one rheometer we used the measuring plate PP20 (part no. 3049) and for the other one the shaft for disposable measuring systems (part no. 10636) with disposable plate D-PP20 (cat. no.17473, Anton Paar). Collagen solutions (with or without cells/hydrogel microparticles) were deposited on the bottom plate, preheated to 37°C. After quickly lowering the top plate to the desired gap, 500 *µ*L of complete medium was pipetted around the sample edge to prevent drying. The time evolution of the storage and loss moduli (*G^′^* and *G^′′^*, respectively) during collagen polymerization was monitored every 5 s by applying a small-amplitude (*γ* = 1%) oscillatory strain at a constant frequency *f* = 1 Hz for 90 minutes until the moduli reached their steady-state values. Next, the frequency-dependent shear moduli of the final network were measured by applying oscillations with a constant strain amplitude (*γ* = 1%) and a frequency ranging from 0.1 Hz to 10 Hz. Finally, the nonlinear elastic response and rupture stress of the network was tested by applying a stress ramp, with stresses logarithmically increasing from 0.01 Pa to 100 Pa at a rate of 10 points per decade and with 5 s between each point.

### 2.6. Flow cytometry measurements of cell sizes

The size distributions of MDA-MB-231 Lifeact-GFP and HDF cells were evaluated using a Cytek^®^ Amnis^®^ ImageStream^®^X Mk II Imaging Flow Cytometer. Cells were trypsinized (Trypsin-EDTA 0.25%, phenol red, Gibco), centrifuged at 200 g and then resuspended in PBS 1x at a concentration of a few tens of million of cells per mL. The cell diameter was measured using the bright-field channel. Cellular objects taken into account for the measurements had aspect ratios between 0.7 and 1 and areas between 300 and 1400 *µm*^2^ or 150 and 600 *µm*^2^ for HDF or MDA-MB-231 Lifeact-GFP cells, respectively. These thresholds were determined by visual inspection of the images to include only single cells. The average cell diameters over the entire population (6,223 cells for HDF and 13,234 cells for MDA-MB-231 Lifeact-GFP) were used to determine the collagen hydrogel formulation required to compensate for the volume occupied by the cells.

### 2.7. Confocal imaging of collagen networks and collagen degradation

Confocal imaging was performed on collagen networks formed in ‘static conditions’, i.e., outside the rheometer parallel plates. To this end, the networks were polymerized in 18-well glass-bottom plates (81817, Ibidi) at 37°C (Thermomixer Comfort incubator, Eppendorf) in a humid atmosphere for 90 minutes. Complete cell medium was added on top of the gels after 45 minutes to avoid sample drying. The final network structure was visualized using a Zeiss LSM 710 confocal microscope (Carl Zeiss MicroImaging) equipped with an AxioCam MRm camera and a 63x oil objective (Plan-APOCHROMAT 63/1.4 oil DIC ∞/0.17). The collagen network was imaged in reflection with a 514 nm laser. MDA-MB-231 cells expressed Lifeact-GFP and HDF cells were stained with SiR-Actin (Spirochrome) (incubation with 2 *µ*M SiR-actin for 45 minutes at 37°C, 5 % CO_2_). Actin was imaged by fluorescence microscopy with *λ_ex_* = 488 nm and *λ_em_* = 516 nm for Lifeact-GFP and *λ_ex_* = 633 nm and *λ_em_* = 646 nm for SiR-actin. Confocal Z-stacks were acquired over a total depth of 50 *µ*m starting at a height of 10 *µ*m above the coverslip with Z-steps of either 0.35 *µ*m or 1 *µ*m (xy-pixel size = 0.11 *µ*m x 0.11 *µ*m and pixel dwell time 0.78 *µ*s).

Proteolytic degradation of the collagen gels by the cells was tested by confocal imaging of networks labelled with a fluorescent dye-quenched protein substrate (DQ-collagen I). Upon proteolytic cleavage, fluorescence is released and reflects the level of proteolysis by the cells [31]. Lyophilized DQ^TM^ collagen (type I from bovine skin, fluorescein conjugate, D12060, Thermofischer Scientific) was dissolved in sterile milliQ water at a stock concentration of 1 mg/mL. It was mixed at a concentration of 25 *µ*g/mL with the 4 mg/mL collagen preparations. MDA-MB-231 stably transfected with nls-mCherry were embedded at a volume fraction of 4%. The cell-embedded network was polymerized in a humid atmosphere at 37°C using a Thermomixer Comfort incubator (Eppendorf) in 18-well glass-bottom plates (81817, Ibidi) for 90 minutes. Complete medium was added on top of the hydrogels after 45 minutes. The cell-embedded hydrogels were then incubated for 45 minutes at 37°C, 5% CO_2_. Fluorescence resulting from collagen degradation was evaluated with the Zeiss LSM710 confocal microscope. The collagen network was imaged using reflection microscopy with a 514 nm laser and the degradation-induced fluorescence was imaged with a 488 nm laser (*λ_ex_* = 488 nm and *λ_em_* = 516 nm).

### 2.8. Analysis of collagen fiber orientations

To quantify the relative orientation of collagen fibers with respect to the edges of individual hydrogel microparticles and MDA-MB-231 cells (hereafter referred to as “cells”), we employed a custom MATLAB 2021b algorithm taking input from automated analysis using the OrientationJ [32] plugin in Fiji (ImageJ v1.54f) [33]. First, multiple regions of interest (ROIs) were extracted from Z-stack reflection images (collagen channel) using ImageJ. We then considered the maximum intensity projection of Z-stacks (acquired with a step size of 0.35 *µ*m or 1 *µ*m) over a depth of 2 *µ*m, focusing on the central imaging plane of individual cells or microparticles. Each ROI image projection was independently processed using the OrientationJ plugin in ImageJ to obtain the local pixel-based orientation (*θ_absolute_*) with respect to the x-axis. Subsequently, a binary mask of collagen fibers was created using intensity thresholding (using imbinarize in MATLAB), to eliminate noisy orientation values. Cell centroid positions and radii in the same ROI images were extracted using image thresholding: from the actin (Lifeact-GFP) channel (in the case of cells) or by fitting a circular shape to the boundary of the microparticle. Finally, the absolute fiber orientation angles were transformed into polar coordinates centered at each cell’s/microparticle centroid, providing the relative orientation of fibers with respect to the edge of the cell/microparticle of reference. The analysis pipeline is shown in Figure S2.

### 2.9. Rheo-confocal microscopy experiments

A dynamic shear rheometer measuring head (DSR 502, Anton Paar) was equipped with a custom-built bottom stage, designed to be mounted on the Zeiss LSM 710 confocal microscope, substituting the microscope XY-moving stage. A Peltier unit, connected to a control box (PE 94 Temperature Controller, Linkam), was mounted into the bottom stage using a 3D-printed holder to maintain a fixed temperature of 37°C. The Peltier module included a middle hole for optical access. Details are shown in Figure S3. A glass coverslip of 30 mm diameter (No. 1, ECN 631-1585, VWR) was glued to the Peltier unit with a thin layer of silicon glue (5398, Loctite). The upper plate was the same geometry as the one used for the standard rheometers: parallel plate geometry with a diameter of 20 mm (shaft for disposable measuring systems (part no. 10636) with disposable plate D-PP20 (cat. no. 84855, Anton Paar).

In each experiment, first the zero gap was identified by lowering the upper plate and bringing it in contact with the glass bottom plate, with contact defined as the gap where the normal force was 0.1 *N*. The gap was then set to zero on a Mitutoyo dial indicator, which was subsequently used to measure the gap between the plates. After depositing the collagen solutions on the glass coverslip bottom plate, the upper plate was quickly and manually lowered until reaching the gap of 160 *µ*m. To prevent sample drying and maintain humidity, 500 *µ*L of complete medium was pipetted around the sample, and a metal hood with wet tissues was placed around the measuring head (Figure S3).

Similar to the bulk rheology experiments, collagen network formation was monitored for 90 minutes by measuring the increase of the shear moduli using small amplitude oscillatory shear oscillations (*γ* = 1% and *f* = 1 Hz). At the same time, the collagen network (in reflection mode) and cells (in fluorescence mode) were imaged every 10 *s* or 20 *s* in a confocal plane fixed at a height of 30 *µ*m above the coverslip (xy pixel size = 0.11 *µ*m x 0.11 *µ*m and pixel dwell time of 0.78 *µ*s for each channel). The region of interest was located at a fixed radial distance of 0.7 *R*, with *R* being the radius of the top plate. After network formation, we applied a stress ramp with stresses logarithmically increasing from 0.01 Pa to 100 Pa at a rate of 10 points per decade and with 5 s between each point. We imaged the network every 2 s to 15 s (depending on the image size) at a height of 30 *µ*m above the coverslip (xy-pixel size= 0.11 *µ*m x 0.11 *µ*m and pixel dwell time of 0.78 *µ*s for each channel) until the rupture point, after which no reflection signal from the collagen was detected.

### 2.10. Analysis of rheology data

The rheology data were analyzed using a custom-made script written in Python (version 3.11). All the derivatives were computed using the gradient function from the numpy Python library. The network polymerization onset time was defined from time sweeps (storage modulus *G^′^*(*t*) versus time *t*) acquired during polymerization. It was defined as the intersection between the tangent to the curve’s inflection point and the time axis, with the inflection point determined as the maximum of the derivative of the storage modulus with respect to time, 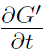 (see Supplementary Figure S4a-b). Before computing the derivatives, the curves were smoothed using the function gaussian filter1d from Python (sigma = 2 for standard rheology and sigma = 10 for rheo-confocal data to compensate for noisier data).

The differential modulus *K* was obtained from the stress-strain curves acquired in stress ramp experiments by calculating the local tangent according to 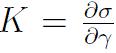. The curves were smoothed using the function gaussian filter1d from Python (sigma = 2). The strain and stress at rupture were identified as the point where *K* reached its maximum value (see Supplementary Figure S4c-d). Plots of the differential modulus as a function of stress were truncated at this rupture point. Curves that showed clear evidence of wall slip, indicated by a non-monotonic trend before rupture, were excluded.

### 2.11. Analysis of confocal images

The kinetics of collagen network formation were determined from the rheo-confocal imaging data by determining the total intensity of reflection microscopy images over time using Fiji (ImageJ). Since cells are also visible in reflection images, we used the fluorescence signal from the GFP-LifeAct stained cells to exclude cell regions. The fluorescence images were processed by applying a Gaussian blur with 2 pixels radius (Fiji) followed by a conversion to a binary image using the Mean or Triangle method in Fiji. The resulting image was subsequently subtracted from the reflection images. Similar to the analysis of the polymerization onset time from the rheological measurements, we determined the onset time from the reflection intensity-time (*I*(*t*)) curves by finding the intersection between the tangent to the curve’s inflection point (point where 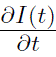 is maximum) and the time axis.

### 2.12. Statistical analysis

For each experiment, the figure captions specify the total number of samples (*n*) and the number of independent experiments (*N*). For boxplots, the center solid line represents the sample median, the box edges correspond to the first and third quartiles and the whiskers are equal to 1.5 times the inter-quartile range. All statistical tests were carried out with Mann-Whitney tests using the scipy Python library. Statistical significance is indicated using p-values (*, ** and *** corresponding to p*<*0.05, p*<*0.01 and p*<*0.001, respectively) and *ns* denotes non-significant differences.

## 3. Results

### 3.1. Cancer cells and fibroblasts have opposite effects on collagen network mechanics

To investigate how different human cells influence the bulk mechanics of collagen networks, we first compared the effect of highly invasive cancer cells (MDA-MB-231) and human dermal fibroblasts (HDFs). Cells were mixed into 4 mg/mL collagen solutions at various volume fractions, ranging from 0.4% to 20%. This collagen concentration replicates the collagen density of breast tumor microenvironments [34]. To ensure a constant collagen network density, we compensated for the presence of the cells by adjusting the hydrogel formulation to account for the volume occupied by the cells calculated from their average diameter (29.3 ± 4.2 *µ*m for HDF cells and 19.7 ± 2.1 *µ*m for MDA-MB-231 cells, see Figure S1). The cell-collagen mixture was polymerized between the two parallel plates of a shear rheometer with the plates spaced apart by 160 *µ*m (see schematic in Figure 1a left). Collagen polymerization was monitored over a time period of 90 minutes by applying small-amplitude strain oscillations to measure the storage modulus *G^′^* and loss modulus *G^′′^* (Figure 1a middle). Next, the final network was subjected to a stress ramp to assess the nonlinear elastic response and rupture strength (Figure 1a right). The choice of gap size did not affect the mechanical properties of the networks (Figure S5), indicating that no wall slip occurred during the measurements. Similarly, the application of a small oscillatory strain during polymerization had no effect on the network mechanics (Figure S6).

**Figure 1:**
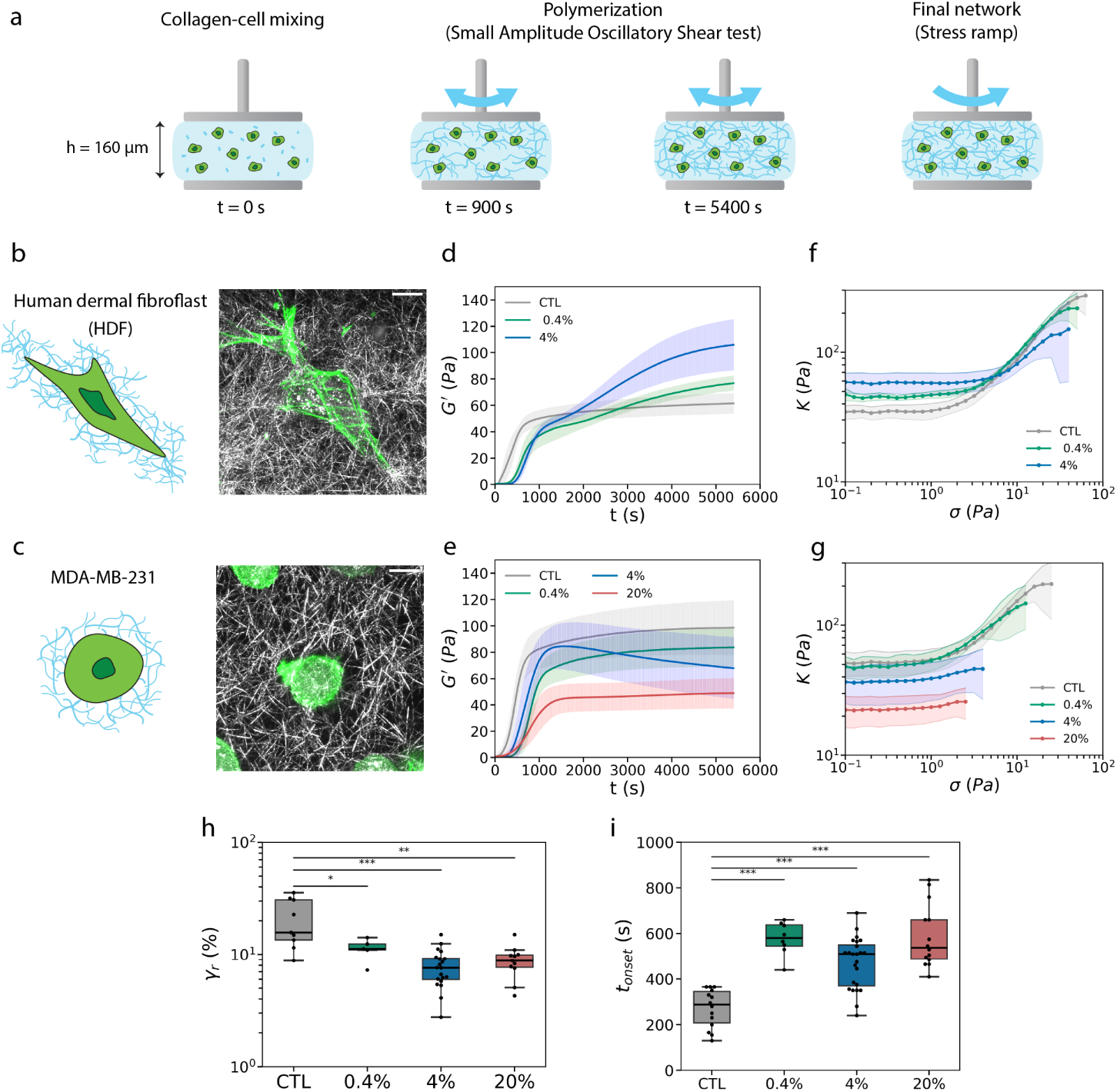
Invasive breast cancer cells (MDA-MB-231) and human dermal fibroblasts (HDF) have opposite effects on the bulk rheology of collagen networks. (a) Cells were mixed with a solution of type I collagen monomers and confined between the two parallel plates of a shear rheometer at t=0 (left). Collagen polymerization was monitored by applying small-amplitude shear oscillations (middle). The final network was subjected to a stress ramp to assess the nonlinear elastic response and rupture strength (right). (b-c) Fibroblasts (b) and breast cancer cells (c) embedded in a collagen network. Schematics (left) and maximum intensity projections of confocal Z-stacks (right). Collagen fibers (grey) were imaged using reflection and the cellular actin cytoskeleton (green) was imaged via fluorescence (using SiR-actin labelling for HDF cells and Lifeact-GFP for MDA-MB-231 cells). Scale bars are 10 *µ*m. (d-e) Storage modulus *G*^′^ as a function of polymerization time for various volume fractions of HDF cells (d, CTL: n=4, N=1; 0.4%: n=2, N=1; 4%: n=4, N=1) and MDA-MB-231 cells (e, CTL: n=16, N=8; 0.4%: n=8, N=3; 4%: n=24, N=10; 20%: n=16, N=8). Data represent mean ± SD. (f-g) Differential modulus *K* as a function of applied shear stress for cell-embedded collagen networks with various volume fractions of HDF (f, CTL: n=3, N=1; 0.4%: n=2, N=1; 4%: n=4, N=1) and MDA-MB-231 cells (g, CTL: n=9, N=4; 0.4%: n=5, N=2; 4%: n=20, N=8; 20%: n=11, N=6). Data represent mean ± SD. (h-i) Boxplots of the strain at rupture *γ_r_* (h) and the collagen polymerization onset time *t_onset_* (i) measured for control networks (CTL) and networks containing MDA-MB-231 cells at volume fractions of 0.4%, 4% or 20%.

Confocal fluorescence imaging of the actin cytoskeleton of the cells, once collagen polymerization was complete, showed that the fibroblasts were elongated with branched protrusions extending from the cell poles (green signal in Figure 1b), whereas the cancer cells were roundish and showed only small protrusions (Figure 1c). This difference in cell morphologies suggests that fibroblasts interact more persistently with the collagen matrix and may exert greater long-term contractile forces on the network compared to cancer cells.

This was further confirmed by reflectance imaging of the collagen network surrounding the cells, that showed accumulation of collagen fibers (grey signal in Figure 1b) near the fibroblasts protrusions, but a more homogeneous collagen fiber distribution around the cancer cells (Figure 1c).

Time-resolved rheology experiments showed that these morphological differences were associated with opposite effects of the cells on the bulk mechanics of the collagen network: the fibroblasts caused global stiffening of the collagen matrix, whereas the cancer cells caused a global softening, relative to control networks (Figure 1d). For control networks (in absence of cells), the storage modulus *G^′^* suddenly started to increase after a delay time of a few hundred seconds, and then quickly reached a constant value. This kinetic behavior reflects the known nucleation-and-growth mechanism of collagen polymerization [35, 36]. In the presence of fibroblasts, the storage modulus showed a biphasic increase with time. The first stiffening phase set in at a time point that was only weakly dependent on cell density, suggesting that it originates from collagen fiber nucleation (Figure S7). The second phase set in at a time point that did depend on cell volume fraction (compare 0.4% and 4%, individual curves in Figure S8), suggesting that it is due to cell-mediated contraction. The loss modulus *G^′′^* showed similar time dependencies as the storage modulus (Figure S9). In marked contrast to the fibroblasts, the invasive cancer cells (MDA-MB-231) caused global softening of the collagen network relative to control networks (Figure 1e). This effect was dependent on cell density. Cells at 0.4% did not significantly impact the final storage modulus. Strikingly, however, cells at 4% caused a non-monotonic time-dependent modulus. The elastic modulus first increased, peaking at a value similar to the control network, and then gradually decreased to a final value about two-fold lower than for the control networks. This non-monotonic behavior was observed for more than 75% of the samples (Figure S10). When the cell volume fraction was further raised to 20%, the non-monotonic behavior was lost, but the final network was again about two-fold softer than the control network. We did not observe any obvious changes in the collagen fiber length or density in the network surrounding the cells to explain the softening (Figure S11). We note that the loss modulus *G^′′^*showed similar time dependencies as the storage modulus (Figure S12). Overall, the only similarity between the behavior of the fibroblasts and MDA-MB-231 cells was that they both delayed the onset of collagen polymerization (Figure 1i).

Following collagen polymerization, the nonlinear elastic behavior of the mature network was assessed by applying a gradually increasing shear stress until network rupture. We quantified the nonlinear response via the differential modulus *K*, defined as the local derivative of the stress/strain curves (*K* = *<u>^∂σ^</u>*). With increasing stress, control networks showed an initial linear response (constant *K* values), followed by stress-stiffening above stresses of ∼ 1 Pa (grey curves in Figure 1f and g). Stress-stiffening behavior is a wellknown feature of collagen networks that is attributed to a transition from a soft-bending-dominated regime at small deformations to a rigid stretch-dominated regime at large deformations [14, 37, 38]. Fibroblasts and cancer cells had very different effects on this stress-stiffening behavior. Fibroblasts only minimally affected the stress-stiffening behavior of collagen, irrespective of cell volume fraction (Figure 1f, individual curves in Figure S13a-c). Furthermore these cells did not affect the rupture strain nor the rupture stress (Figure S13d-e). By contrast, the cancer cells strongly inhibited the stress-stiffening behavior of collagen above a cell volume fraction of 4% (Figure 1g). They also strongly reduced the rupture strain *γ_r_* (Figure 1h) and rupture stress *σ_r_* (Figure S14) compared to the control collagen network. We conclude that fibroblasts and cancer cells have opposite effects on the bulk rheology of collagen. Fibroblasts apply contractile stress and cause network stiffening, whereas the cancer cells apply minimal contractile stress, cause network softening, and suppress stress-stiffening.

### 3.2. Both volume exclusion and cell adhesion impact collagen network mechanics

The embedded cancer cells could modulate collagen network mechanics by acting as viscoelastic inclusions [15] and/or by actively exerting forces on the collagen fibers through integrin-mediated adhesion receptors [39]. To test the importance of volume exclusion, we first performed bulk rheological measurements on collagen networks with embedded soft hydrogel microparticles. These microparticles are inert, cell-sized, mechanically uniform and isotropic [27]. Importantly, they lack collagen-binding sites on their surface, so they merely serve as soft passive cell-sized inclusions [40, 41]. Confocal imaging showed that collagen fibers in close proximity of the microparticles (MPs) were oriented tangentially to their surface (Figure 2a and Figure S15a-b), in contrast to the isotropic orientation of collagen fibers around the cancer cells (Figure 1c, Figure S15a-b). Nevertheless, the passive microparticles caused a similar reduction in the elastic modulus of the collagen networks (Figure 2b) as the cancer cells. The microparticles also similarly delayed collagen polymerization (Figure 2h), indicating that passive inclusions slow down the collagen polymerization process. At 20% volume fraction, the inclusion of microparticles resulted in a two-fold reduction in the network stiffness, comparable to the decrease observed with cancer cells. This result suggests that the softening observed at higher cell volume fractions is driven by a volume exclusion effect rather than cell-matrix adhesion.

**Figure 2:**
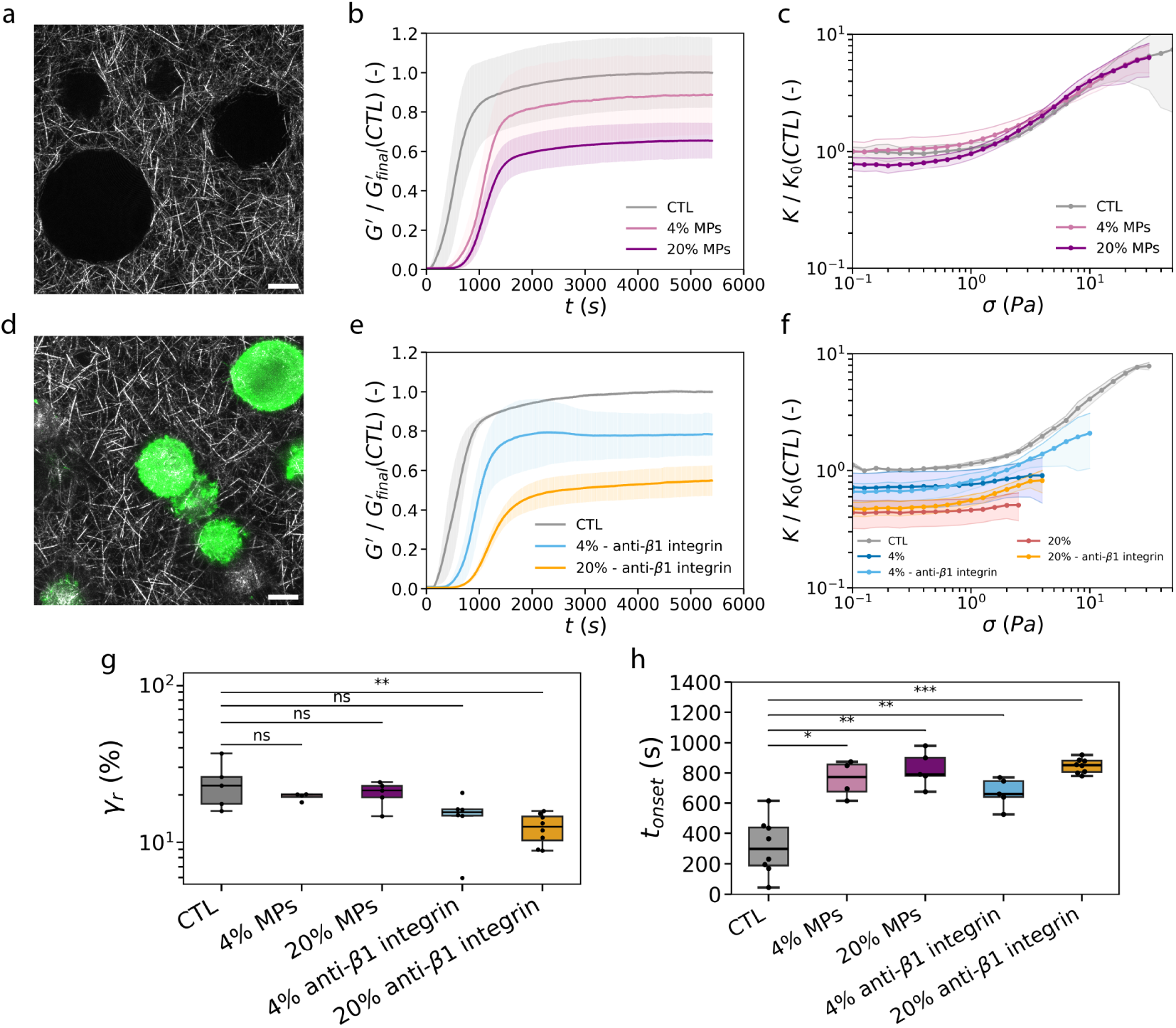
Volume exclusion and cell adhesion together determine the impact of cancer cells on the bulk mechanics of collagen networks. (a,d) Maximum intensity projections of Z-stacks of confocal images across a depth of 10 *µm* of a collagen network with embedded soft hydrogel microparticles (a) or MDA-MB-231 cancer cells with an adhesion-blocking anti-*β*1 integrin antibody (d). Scale bars are 10 *µm*. Collagen (grey) is imaged in reflection and the cells (green) in fluorescence (using LifeAct-GFP labeling). The dark circles in (a) are due to the presence of microparticles. (b,e) Storage modulus *G*^′^ normalized to the final modulus of control collagen (*G*^′^ (*CTL*)) as a function of polymerization time for networks with hydrogel microparticles (MPs) (b, CTL: n=6, N=2; 4%: n=4, N=2; 20%: n=5, N=2) or adhesion-blocked MDA-MB-231 cells (e, CTL: n=2, N=1; 4%: n=5, N=2; 20%: n=8, N=2), at volume fractions of 0% (CTL), 4% and 20%. Data represent mean ± SD. (c,f) Differential modulus *K* normalized to the linear modulus of control collagen (*K*_0_ (*CTL*)) as a function of applied shear stress *σ* for networks containing hydrogel microparticles (c, CTL: n=3, N=2; 4%: n=4, N=2; 20%: n=5, N=2) or adhesion-blocked MDA-MB-231 cells (f, CTL: n=2, N=1; 4%: n=5, N=2; 20%: n=8, N=2) at volume fractions of 0% (CTL), 4% and 20%. (g-h) Boxplots of the strain at rupture *γ_r_* (g) and of the collagen polymerization onset time *t_onset_* (h) for collagen networks containing hydrogel microparticles or adhesion-inhibited MDA-MB-231 cells at volume fractions of 0% (CTL), 4% and 20%.

At the lower volume fraction of 4%, however, we observed an interesting difference in the time dependence of the modulus for microparticles versus cells: the time dependence was monotonic for microparticles (Figure 2b), whereas it was non-monotonic for cancer cells (Figure 1e). Stress ramp experiments revealed another notable difference: while the cancer cells suppress stress-stiffening, the hydrogel microparticles preserved the stress-stiffening behavior of collagen networks across all volume fractions (4% and 20%) (Figure 2c). Stress-stiffening curves for networks with microparticles normalized to the linear modulus of the corresponding control network (*K*_0_(*CTL*)) overlapped with the control collagen at stresses above ∼ 2 Pa. The microparticles also did not significantly change the network rupture strain (Figure 2g) nor rupture stress (Figure S16), in contrast to the reduced strength in presence of the cancer cells.

These observations suggest that volume exclusion only partially explains the impact of cancer cells on collagen network mechanics. To test whether integrin-mediated adhesion also plays a role, we embedded cancer cells in collagen while blocking *β*1-integrin adhesion receptors with a specific antibody. *β*1-integrins are known to mediate cell adhesion to collagen type I fibers [42]. Confocal imaging showed that collagen fibers in close proximity of cancer cells with blocked integrins were more tangentially oriented as compared to fibers around untreated cells (Figure 2d and Figure S15a-b), although this effect was less pronounced than the tangential orientation observed around the hydrogel microparticles. Further away from the adhesion-inhibited cells and the microparticles (beyond 4 *µ*m), the collagen fibers were instead arranged isotropically (Figure S15c). Similar to the microparticles, the adhesion-inhibited cells delayed the onset of collagen polymerization (Figure 2h) and reduced the final network modulus (Figure 2e, for individual curves see Figure S17). At a volume fraction of 20%, the adhesion-inhibited cells reduced the modulus by about two-fold compared to control networks, comparable to the impact of the non-adhesive microparticles. At a volume fraction of 4%, collagen with adhesion-inhibited cells showed a monotonic increase of the modulus with time, similar to collagen with microparticles. Finally, collagen networks with adhesion-inhibited cells showed stress-stiffening across all cell volume fractions (Figure 2f), although less marked than with microparticles.

There were striking differences between collagen gels with adhesion-inhibited cells compared to cells capable of integrin-mediated adhesion. Networks with cells capable of adhesion showed a non-monotonic dependence of the storage modulus with time during collagen polymerization, not seen upon integrin blocking. Also, networks with adherent cells did not stress-stiffen and had smaller rupture strains (Figure 1h) and stresses (Figure S16) than networks with adhesion-inhibited cells (Figure 2g). These observations suggest that the cancer cells affect collagen mechanics by a combination of volume exclusion and adhesion-dependent effects. Non-adherent particles and cells (irrespective of adhesion) cause a delay in collagen polymerization and lowering of the final network modulus at high enough volume fraction of the inclusions. These are apparently effects caused by volume exclusion. However, the time-dependent decrease in the storage modulus at low cell volume fraction (4%) and the loss of collagen stress-stiffening are directly associated with the cells’ ability to adhere to collagen fibers.

### 3.3. Rheo-confocal imaging reveals dynamic cell-mediated network remodeling

To understand why cancer cell adhesion impacts the bulk mechanics of the collagen networks, we integrated a rotational rheometer with an inverted confocal microscope, enabling simultaneous rheological measurements and confocal imaging of the collagen network and embedded cells (Figure 3a). Briefly, a shear rheometer measuring head was placed on top of a confocal microscope, substituting its XY-moving stage (Figure S3). A glass cover-slip was used as bottom plate, to allow high resolution confocal imaging of the samples from below (with minimal effect on the apparent rheological measurements as shown in Figure S18).

**Figure 3:**
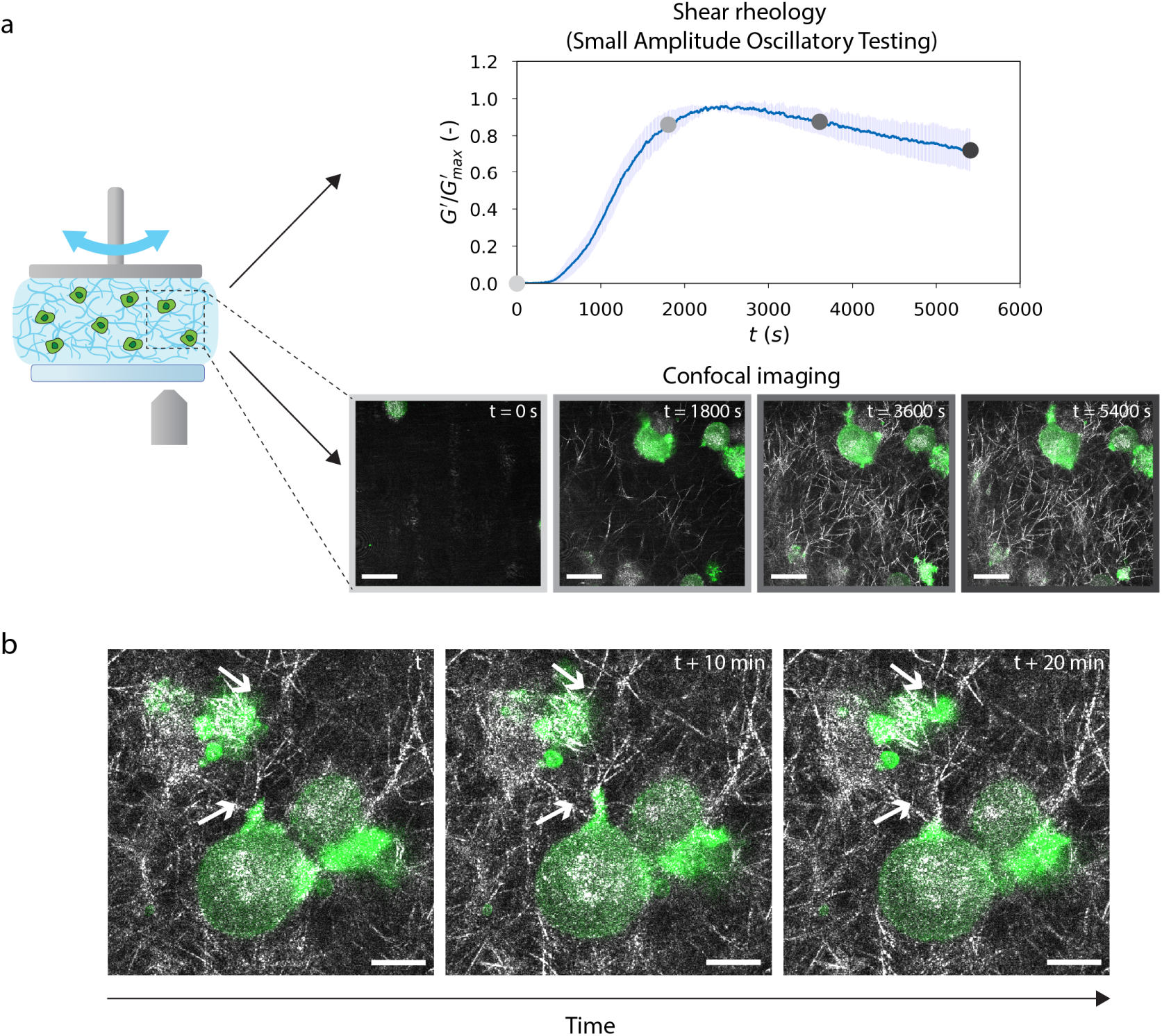
Rheo-confocal microscopy demonstrates dynamic local remodelling of the collagen network by the cancer cells via actin-mediated protrusions. (a) Schematic of the rheo-confocal setup allowing simultaneous shear rheology measurements and confocal imaging. Top row: Storage modulus *G*^′^ normalized by its maximum *G*^′^ value as a function of polymerization time for a collagen network containing 4% MDA-MB-231 cells. Data represent mean ± SD (*N* = 3). Bottom row: Corresponding representative confocal images at different time points (indicated by colored circles in the shear rheology curve). Collagen fibers are visible in grey (reflection microscopy) and actin labeled with LifeAct-GFP (fluorescence microscopy) is shown in green. Scale bars are 20 *µm*. (b) Representative time-lapse confocal images for a collagen network (grey) containing 4% MDA-MB-231 cells (green) at different times after the initiation of polymerization. We observe cell-mediated collagen fiber bending (white arrow at the top of the images) and fiber pulling (white arrow at the center of the images). These local remodeling events occur for collagen fibers associated with small cell protrusions. Scale bars are 10 *µm*.

We first used the rheo-confocal setup to correlate the time evolution of the shear moduli during collagen polymerization with the underlying changes in network structure. Confocal imaging showed that collagen fibers first appeared around the same time that the storage modulus started to suddenly increase (Figure S19a). This was observed both in the absence (Supplementary Video 1) and presence (Supplementary Video 2) of cancer cells. The onset times determined from the time-dependent increase of the normalized intensity of the collagen networks (reflection channel) were longer than those obtained from the storage modulus (Figure S19b-d). This discrepancy may result from the inability to detect fibers below a certain thickness by reflection microscopy [43] and the limitation of imaging to a specific region of the sample. Importantly, both confocal imaging and rheology showed that cancer cells increased the onset times for collagen polymerization by about 50% as compared to control collagen networks, confirming that the presence of cells delays nucleation and growth of collagen fibers.

Confocal fluorescence imaging showed that the cancer cells actively remodeled the collagen network in a dynamic way as the network formed (Supplementary Videos 3-5). The cancer cells extended and retracted actin-containing membrane protrusions, which mechanically engaged with collagen fibers. The high F-actin intensity in cell protrusions coincided with regions where the collagen networks was visibly remodeled (Figure 3b and Supplementary Videos 3-5), indicating that the cells remodel the network through active force application. We observed clear examples of collagen fiber bending (see example denoted by the arrow at the top of the time lapse image series in Figure 3b) and pulling (see example denoted by the arrow in the center of the time lapse image series in Figure 3b). These fiber deformations were transient, relaxing after the cells retracted the protrusions. Thus, cancer cells can transiently (on timescales of tens of minutes) alter the local structure of the surrounding collagen network through active mechanical interactions with collagen fibers.

Similar imaging of the mature collagen network, with or without embedded cancer cells, was performed during the shear stress ramp (Supplementary Videos 6-7). Upon shear, fibers and cells exhibited co-translation up to the rupture point, indicated by the loss of reflection signal. We could not observe any notable differences in collagen fiber displacement under shear between control networks versus networks with cells to explain the loss of stress-stiffening or the reduced rupture strength. In contrast to actively remodeling cancer cells, we also examined the effect of passive, non-adhesive microparticles (at 20% volume fraction) embedded in the collagen network in the rheo-confocal setup (Supplementary Videos 8–9). During polymerization under oscillatory shear (Supplementary Video 8), collagen fibers appeared in random locations with no evidence of preferential nucleation around MPs, which were visualized as voids in the reflection signal. Under a shear stress ramp (Supplementary Video 9), the MPs and surrounding fibers exhibited global lateral co-translation up to the rupture point, where MPs remained undeformed up to about 10 Pa. This comparison highlights that while cancer cells actively remodel and mechanically engage with the collagen network, passive microparticles primarily influence network behavior in a volume-exclusion manner.

### 3.4. Active cell contractility contributes to collagen network softening

The rheo-confocal experiments revealed that cancer cells actively interact with collagen fibers through actin-filled protrusions and bend, pull and displace collagen fibers. Combined with the observation that blocking integrin-mediated cell adhesion prevented time-dependent softening of cell-embedded gels and restored stress-stiffening, we hypothesized that active myosin-driven contraction may contribute to the mechanics of cell-embedded networks. To test this hypothesis, we inhibited cell contractility with (±)-blebbistatin, a specific myosin II ATPase inhibitor. Since blebbistatin is dissolved in dimethyl sulfoxide (DMSO), we first verified that exposure to DMSO (0.2% volume fraction) did not affect the cells nor their interaction with the collagen network (Figure S20).

Interestingly, the effect of blebbistatin on the rheology of cell-embedded collagen networks depended on cell density. At high cell volume fraction, the polymerization behavior and the final storage modulus and stress-stiffening behavior of the networks (Figure 4a-b) were similar as in absence of blebbistatin. By contrast, blebbistatin significantly changed these features for collagen networks containing cells at an intermediate (4%) volume fraction. First, blebbistatin addition prevented the non-monotonic time-dependence of the storage modulus with a peak at intermediate times seen in its absence (Figure 4c, individual curves in Figure S21). Thus, in presence of blebbistatin, the polymerization curves were comparable to those of control collagen and collagen with adhesion-blocked cells. This finding indicates that both cell adhesion and myosin-based contractility are needed for time-dependent network softening by cancer cells. An alternative explanation for cell-mediated network softening could be proteolytic degradation by cell-secreted or cell-surface enzymes. We could, however, exclude this explanation, as no collagen degradation was observed on our experimental time scale (Figure S22). Second, blebbistatin addition recovered stress-stiffening of cell-embedded networks (Figure 4d), which was impaired in absence of blebbistatin. Also, blebbistatin returned the rupture strain (Figure 4e) and stress (Figure S24) to values close to those of control collagen networks. Apparently, the cancer cells suppress stress-stiffening of collagen networks by physical remodeling, which requires integrin-based adhesion to collagen and myosin-driven active forces.

**Figure 4:**
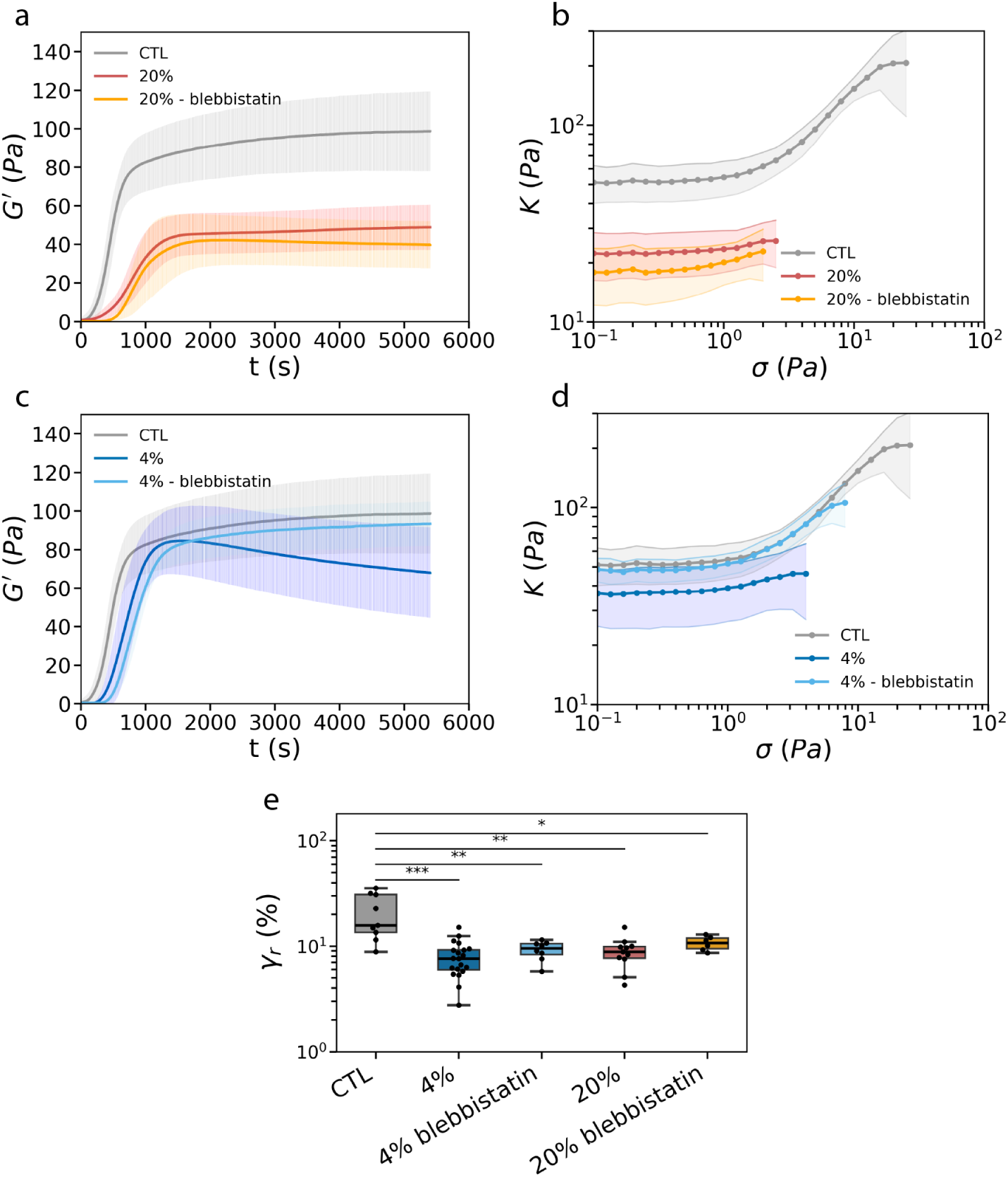
Inhibiting myosin-based contractility of cancer cells demonstrates that cells influence collagen mechanics through active remodeling only at intermediate cell volume fractions. (a,c) Storage modulus *G*^′^ as a function of polymerization time for collagen networks containing MDA-MB-231 cells at a volume fraction of 20% (a, CTL: n=16, N=8; 20%: n=16, N=8; 20%-blebbistatin: n=6, N=2) or 4% (c, CTL: n=16, N=8; 4%: n=24, N=10; 4%-blebbistatin: n=8, N=2), comparing control conditions (no cells) and cell-embedded networks with or without blebbistatin treatment (3 hours at 10 *µ*M). (b,d) Differential modulus *K* as a function of applied shear stress *σ* for collagen networks containing MDA-MB-231 cells at a volume fraction of 20% (b, CTL: n=9, N=4; 20%: n=11, N=6; 20%-blebbistatin: n=6, N=2) or 4% (d, CTL: n=9, N=4; 4%: n=20, N=8; 4%-blebbistatin: n=8, N=2), comparing control conditions (no cells) and cell-embedded networks with or without blebbistatin. Data represent mean ± SD. (e) Boxplots of the strain at rupture (*γ_r_*) for collagen networks with 0%, 4% or 20% cells, with or without blebbistatin treatment.

## 4. Discussion

Cells play a critical role in shaping the mechanical properties of their surrounding matrix [4, 44]. They can influence the matrix through multiple interconnected processes, including volume exclusion effects [45, 46, 15, 16], adhesion [47], mechanical force generation [26], and biochemical modifications [48]. Since these mechanisms often act simultaneously, decoupling each of their effects on the matrix mechanics is challenging. Here, we investigated how cancer cells influence the shear mechanics of collagen type I networks in bulk rheology experiments. To disentangle the contribution of each individual process, we used cell-sized microparticles to test the impact of passive inclusions, myosin and integrin inhibitors to test the influence of cell adhesion and contractility.

Different cell types (cancer cells and fibroblasts), but also passive hydrogel microparticles, delayed the onset of collagen polymerization even at small volume fractions. Rheo-confocal experiments showed via simultaneous imaging and rheology that collagen networks are formed through the nucleation of collagen fibers, which rapidly grow and thicken as the network matures. Both cells and microparticles delayed the first appearance of fibers. Apparently, the space occupied by the inclusions reduces the available space for collagen polymerization and thereby delays the formation of a network-spanning structure, consistent with the theory of gelation through percolation [49]. Interestingly, the effect of cell-sized inclusions is opposite to that of small (100 nm) extracellular vesicles, which were recently shown to accelerate collagen fibrillogenesis [50]. While extracellular vesicles likely act as nucleation points, much larger (tens of micrometers) cells and microparticles delay network formation through a passive volume exclusion effect.

The final stiffness of the collagen networks was strongly dependent on the type of inclusion. Human dermal fibroblasts stiffened the collagen networks even at the lowest volume fraction tested (0.4%) (Figure 1d). By contrast, cancer cells softened the collagen networks at volume fractions between 4% and 20%. Both cell types adhere to and actively interact with the collagen fibers, but their distinct effects on stiffness arise from differences in the nature and timescale of these interactions: fibroblasts apply sustained traction that promotes strain-stiffening [51], whereas cancer cells engage in dynamic, transient remodeling that disrupts sustained fiber loading and softens the network. Fibroblasts continuously pull on the collagen network with their filopodia [52]. As a consequence, they generate a significant contractile prestress that stiffens the entire network because of its inherent nonlinear response. A similar fibroblast-induced stiffening effect was previously demonstrated for fibrin networks that exhibit similar stress-stiffening behavior as collagen [53]. Several studies also evidenced a local stiffening of collagen networks around embedded fibroblasts [26, 54, 55, 56], which was attributed to the alignment and recruitment of collagen fibers through pulling forces. Confocal imaging showed that the MDA-MB-231 cancer cells, as compared to fibroblasts, interact in a much more dynamic and transient manner with collagen. The cells extend small protrusions that can bend, pull and displace collagen fibers. However, the protrusions are transient (minute lifetimes) and the fibers are released upon retraction of the protrusions. As a result, there is much less accumulation of collagen fibers around the cancer cells as compared to the fibroblasts (Figure 1b,c). Previous studies measuring local changes in matrix stiffness near cancer cells reported either a local matrix stiffening [26, 56, 57] or no significant mechanical changes [54]. In contrast, our bulk rheology experiments reveal network softening at a macroscopic scale. This discrepancy between local and global mechanical properties is likely a reflection of the highly heterogeneous structure of collagen networks. Computational models have shown that, due to their low connectivity (with mostly three-fold junctions), collagen networks exhibit a highly heterogeneous (non-affine) response to mechanical loading [58, 59].

So, how can the transient and local interactions of cancer cells with the collagen network cause global network softening? At low (4%) cell volume fraction, the cells caused an intriguing time-dependent softening during collagen polymerization. Importantly, this softening was not caused by any proteolytic degradation of the collagen (Figure S22) and the inhibition of metalloproteinase (MMP) activity with batimastat had no significant effect on the rheological behavior within the timescale of our experiments (Figure S23). Instead, cell-mediated softening was dependent on myosin-driven contractility and integrin-mediated adhesion, as shown by experiments where we blocked myosin II activity or *β*1-integrins. Interestingly, a similar inclusion-mediated network softening was recently observed with thermosensitive hydrogel beads that were made to contract by heating [60]. In this study, softening was hypothesized to arise from collagen fiber buckling. Similarly, the cancer cells might buckle collagen fibers in their vicinity.

At higher cell volume fractions (20%), time-dependent softening of the collagen networks was much less striking than at low volume fraction (Figure S10), but the final network was still softer than the cell-free networks. This observation suggests that the global shear mechanics of cell-embedded networks depends on a combination of local active cell-mediated network remodeling and volume exclusion. We surmise that the volume exclusion effect is dominant at higher (20%) volume fractions, where cancer cells and passive microparticles confer a comparable softening effect. It will be interesting to investigate under which physiological circumstances cells can act as such inclusion bodies. For instance, quiescent stromal cells might act similarly to passive inclusions, modulating mechanics mainly through their physical presence [61]. At small (4%) volume fractions, where the space occupied by the cells is smaller, the influence of volume exclusion is smaller, so the time-dependent softening by active collagen remodeling is more dominant. Cell-mediated softening of mature collagen networks has also previously been reported in the presence of Chinese ovary hamster (CHO) cells [62]. This study attributed the softening to the cells’ ability to recruit collagen around themselves, thereby depleting collagen from the remaining network. For the cancer cells studied here, we can exclude this explanation since confocal imaging revealed a uniform collagen density.

Finally, we found that cells also altered the bulk nonlinear elastic response of collagen network mechanics. Fibroblasts and cancer cells again had opposite effects. While fibroblasts merely slightly decreased the strain and stress at rupture, the cancer cells suppressed collagen stress-stiffening and markedly reduced the strain and stress at rupture. Impairment of stress-stiffening required cell-matrix adhesion: when we blocked *β*1-integrin-mediated adhesion, stress-stiffening was (partially) restored. Similarly, passive microparticles that did not adhere to collagen did not affect the stress-stiffening behavior. We speculate that anchoring of the fibers to the cells and the altered collagen organization in the immediate proximity of the cells, acting as defects, hinder long-range stress transmission in the network, leading to loss of stress-stiffening.

## 5. Conclusions

In this study, we showed that invasive cancer cells soften collagen networks and suppress stress-stiffening. This effect is opposite to the more well-studied effect of fibroblasts, which stiffen collagen networks. We showed that softening is caused by a combination of a passive mechanism (volume exclusion) and active mechanisms (network remodeling mediated by cell contractility and adhesion). The balance between these two contributions depends on the cell volume fraction, with the volume exclusion effect predominating at higher cell densities. By using a biomimetic tissue model system, we were able to shed new light on the mechanical role of living cells in fibrous networks, demonstrating how their nonequilibrium activity can influence the global mechanical response under shear. Our work provides a fundamental basis to understand the biophysical processes by which different types of cells impact tissue mechanics in health and disease.

## 6. Credit authorship contribution statement

**Irène Nagle:** Writing – original draft, Visualization, Validation, Software, Methodology, Investigation, Formal analysis, Data curation, Conceptualization. **Margherita Tavasso:** Writing – original draft, Visualization, Validation, Software, Methodology, Investigation, Formal analysis, Data curation, Conceptualization. **Ankur D. Bordoloi:** Software, Formal Analysis, Data curation, Writing – review. **Iain A.A. Muntz:** Software, Methodology, Conceptualization, Writing – review. **Gijsje H. Koenderink:** Writing – review & editing, Validation, Supervision, Project administration, Funding acquisition, Conceptualization. **Pouyan E. Boukany:** Writing – review & editing, Validation, Supervision, Project administration, Funding acquisition, Conceptualization.

## 8. Data and code availability

All data reported in this paper are available upon request. The original codes have been deposited on GitHub and can be accessed publicly via these links: https://github.com/irenenagle/functions_rheology and https://github.com/aerials00/fiber_analysis

## 9. Acknowledgements

I. Nagle and G. H. Koenderink gratefully acknowledge the Flagship Healthy Joints, which is (partly) financed by Convergence Health and Technology, for funding. M. Tavasso and P.E. Boukany gratefully acknowledge funding from the European Research Council (ERC) under the European Union’s Horizon 2020 research and innovation program (grant agreement no. 819424). A.D. Bordoloi gratefully acknowledges funding from MSCA Postdoctoral Fellowships 2022 Project ID: 101111247. MDA-MB-231 Lifeact-GFP and nls-mCherry cells were kindly provided by P. ten Dijke (LUMC Leiden, NL). We thank I. Liang for help with fibroblast cell culture, K. David for help with MDA-MB-231 cell culture and F. Ramirez Gomez for help with the flow cytometry measurements. We thank S. ten Hagen for designing and printing the Peltier holder for the rheo-confocal setup. Flow cytometry measurements for HDF cells were kindly provided by I. Liang and J. Conboy. We thank C. Margadant (Leiden University) for his advice on the usage of the anti-integrin antibody.

## Supplementary Information

### Supplementary information for

**Figure S1:**
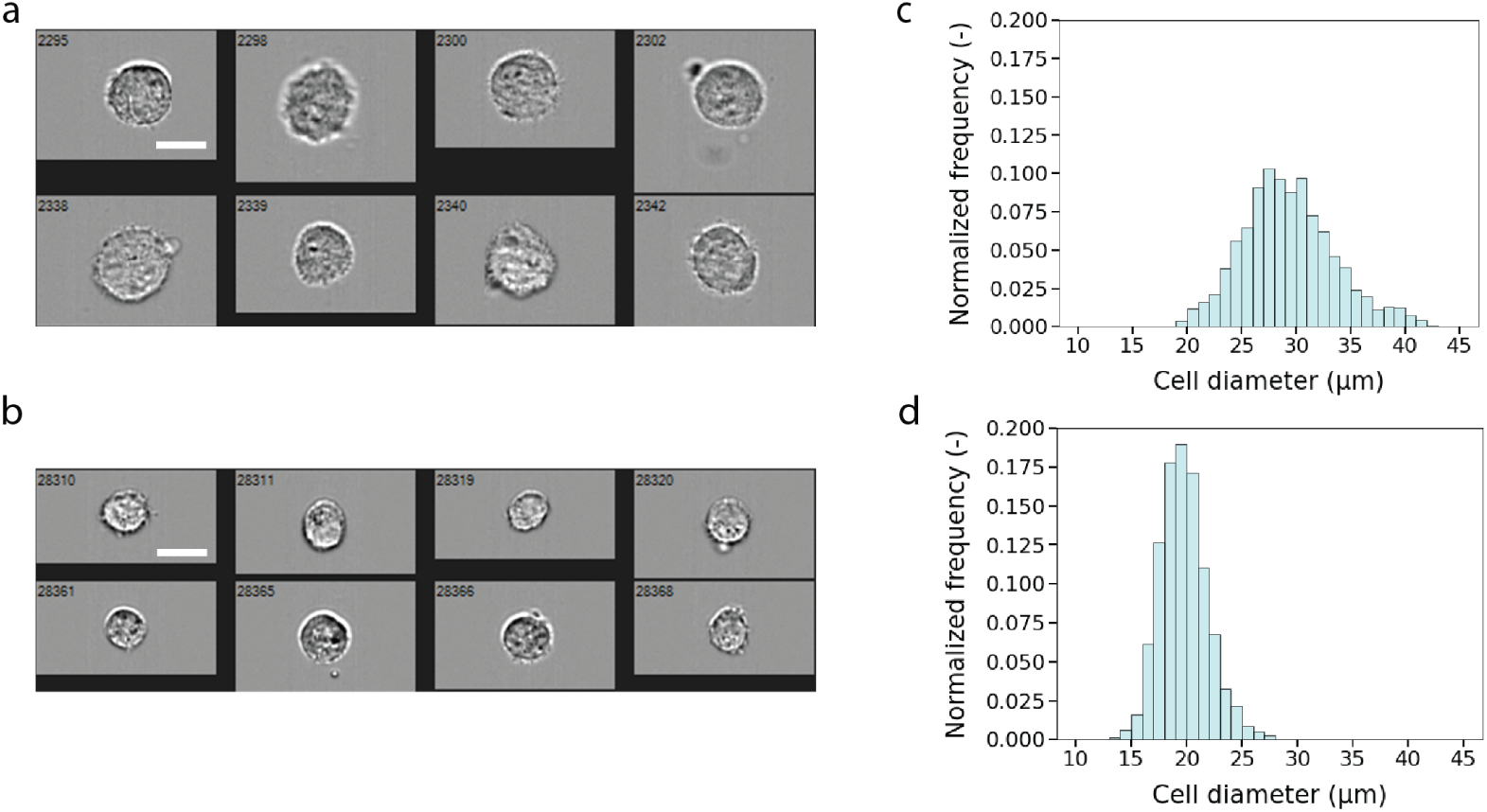
Cell diameter measurements for human dermal fibroblasts (HDF) and breast cancer cells (MDA-MB-231 Lifeact-GFP). (a-b) Representative brightfield images of the HDF cells (a) and MDA-MB-231 Lifeact-GFP cells (b), both imaged in the flow cytometer. Scale bars are 20 *µ*m. (c-d) Cell diameter distributions for HDF cells (c) and MDA-MB-231 Lifeact-GFP cells (d) measured by flow cytometry. The average cell diameter was 29.3 ± 4.2 *µ*m for HDF cells (measured on 6,223 cells) and 19.7 ± 2.1 *µ*m (mean ± SD) for MDA-MB-231 cells (measured on 13,234 cells).

**Figure S2:**
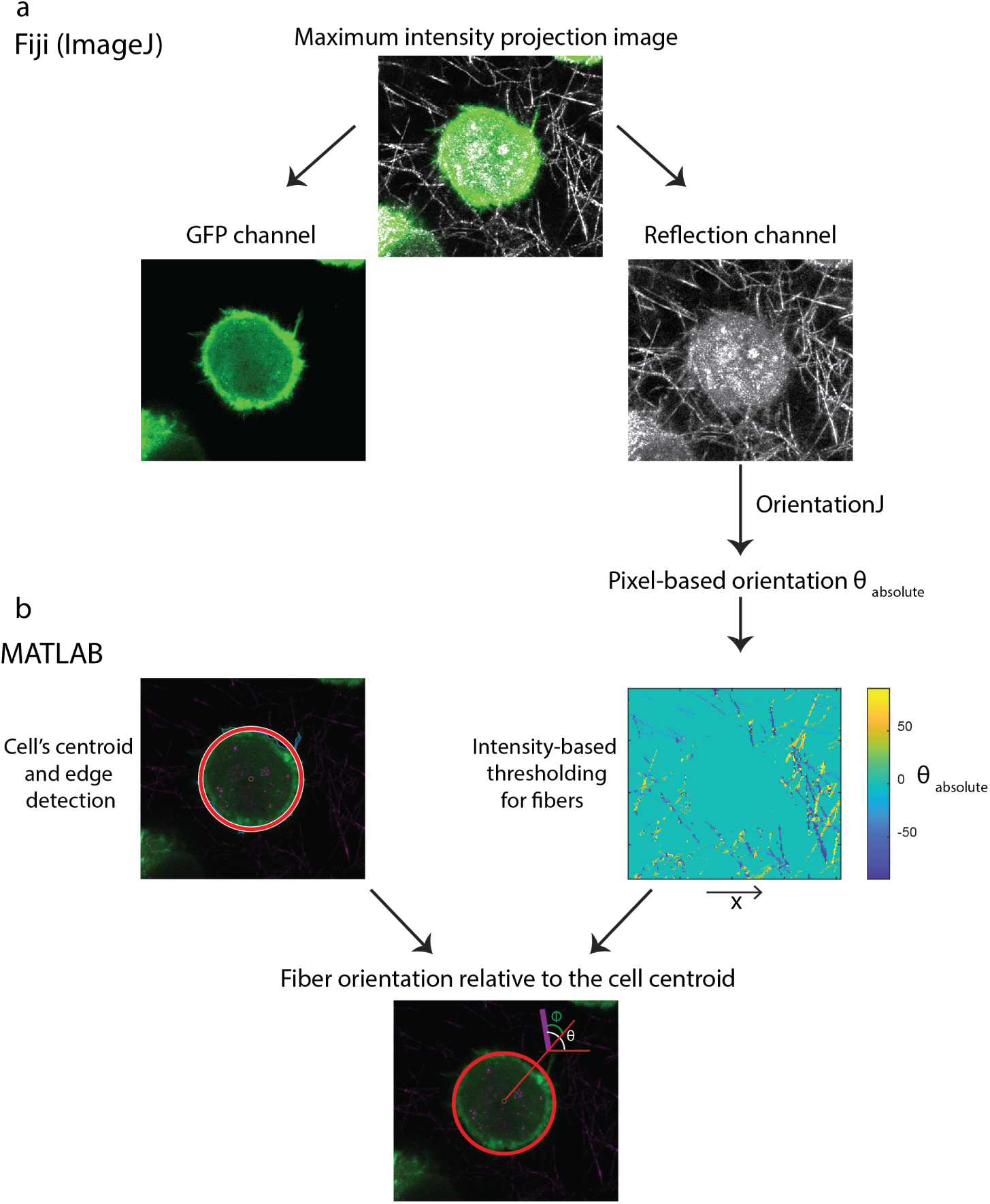
Schematic representation of the analysis pipeline for quantifying collagen fiber orientations relative to microparticles or cells. (a) In ImageJ, a maximum intensity projection over a depth of 2 *µ*m was obtained from Z-stacks. The GFP (actin) and the reflection channels were separated, and the OrientationJ plugin was applied to the reflection image (using the gaussian gradient and a local window *σ* of 1 pixel) to obtain pixel-based orientation angles (*θ_absolute_*) relative to the x-axis. (b) Left: Circular masks are generated based on the cell boundary (red) to define the reference centroid for orientation analysis. For microparticles, a circular shape was fitted to the boundary to create a binary mask, as they are not fluorescent. Right: The pixel locations corresponding to the cell/microparticle were subtracted from the OrientationJ output in order to process only pixels from collagen fibers. An intensity-based thresholding was applied (using the function imbinarize in MATLAB), to remove noisy orientation values. The colormap represents the extracted orientation values (*θ*_absolute_) of the selected fibers. Bottom: The absolute orientation angles (*θ*_absolute_, white) were converted into polar coordinates centered at the cell/microparticle centroid to determine the relative orientation *ϕ* (green) for each fiber pixel (example fiber in purple).

**Figure S3:**
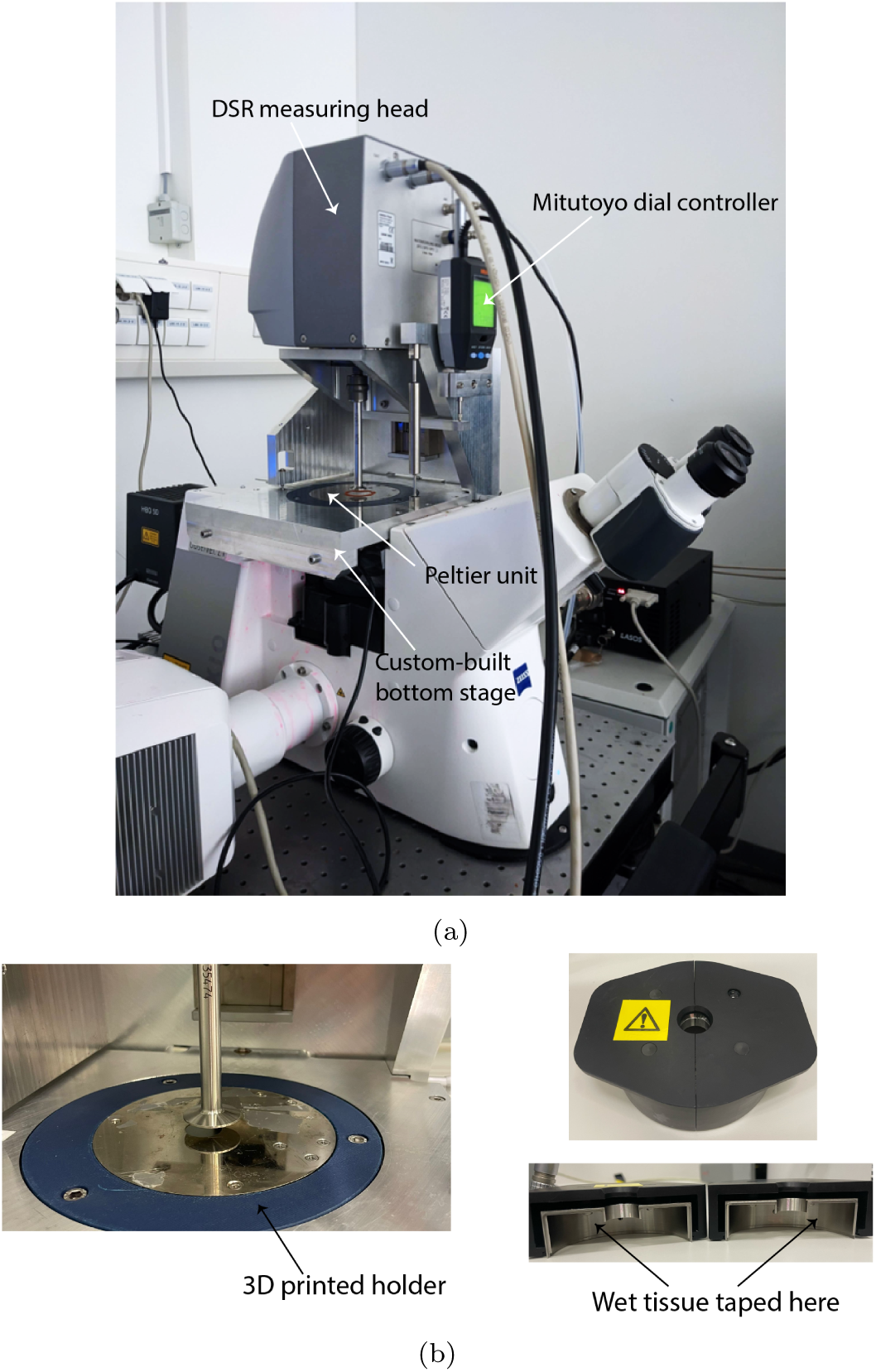
Rheo-confocal microscopy setup. (a) Photograph of the setup shows the DSR rheometer measuring head mounted on a confocal microscope through a custom-built bottom stage that replaces the microscope’s original XY-moving stage. The measuring gap is controlled by a Mitutoyo dial indicator, while the temperature is controlled by a Peltier unit, on which a glass coverslip is glued (red circle in the image) for sample deposition and imaging. (b) Left: The Peltier unit was mounted onto the rheometer bottom stage using a 3D-printed holder, made of polylactide (PLA). Right: A metal hood with wet tissues taped inside was placed around the measuring head to maintain humidity and prevent sample drying.

**Figure S4:**
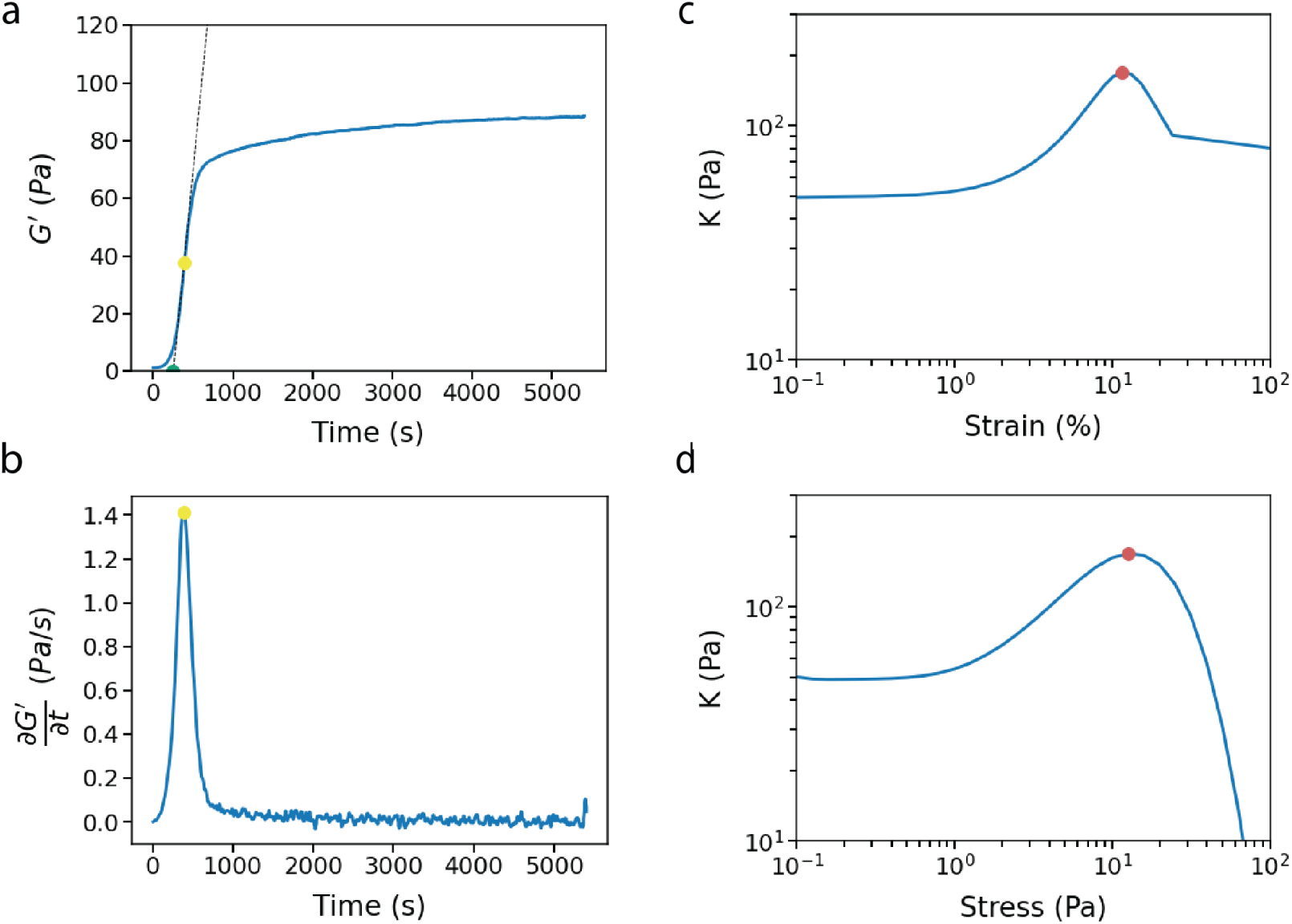
Determination of the onset time and strain/stress at rupture. (a-b) Representative curves of the storage modulus *G*^′^ (a) and the derivative of the storage modulus with respect to time 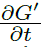 (b) as a function of time. The onset time (green point) is determined from the polymerization curve (a) as the intersection between the tangent (dashed black line) to the curve’s inflection point (yellow point) and the time axis. The inflection point of the curve is determined as the maximum of 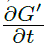. (c-d) Representative curves of the differential modulus *K* as a function of the strain *γ* (c) or the stress *σ* (d). The strain and stress at rupture are determined as the point where the differential modulus reaches its maximum value (red point).

**Figure S5:**
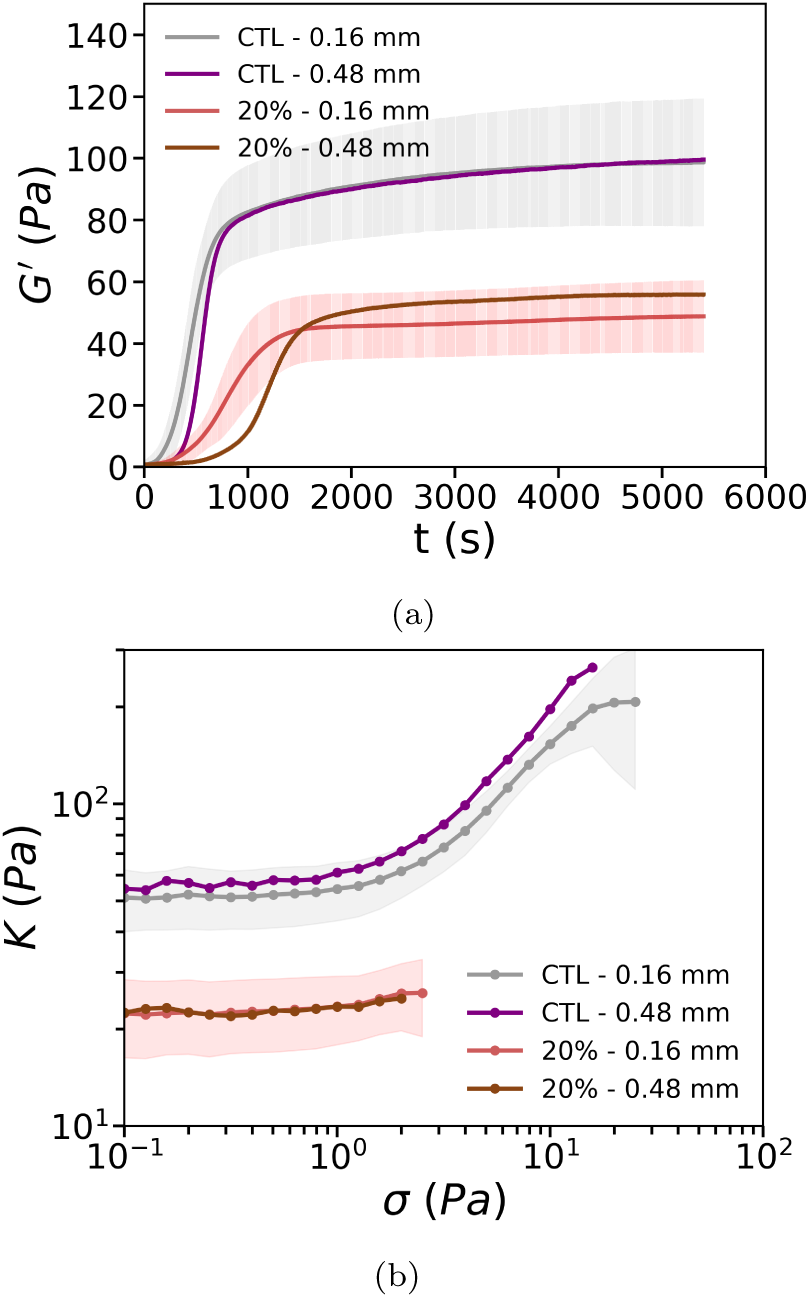
The measurement gap of the parallel plate geometry does not influence the bulk rheological properties of collagen networks. (a) Time-dependent storage moduli *G*^′^(*t*) during polymerization for control collagen networks and cancer cell-embedded networks (at 20% volume fraction of MDA-MB-231 cells) for two different gaps (0.16 mm and 0.48 mm). The longer delay before the onset of collagen polymerization for control networks and networks containing 20% cells observed when the gap is 0.48 mm gap is likely due to the larger sample volume deposited between the plates. (b) Corresponding stress-dependent differential moduli *K*(*σ*) measured by applying a stress ramp to the mature networks. Curves are truncated at the rupture point. Comparable results are obtained for gaps of 0.16 mm (CTL: n=16, N=8 and 20%: n=16, N=8) and 0.48 mm (CTL and 20%: n=1, N=1), confirming that wall slip effects are negligible. Graphs in (a) and (b) represent mean ± SD.

**Figure S6:**
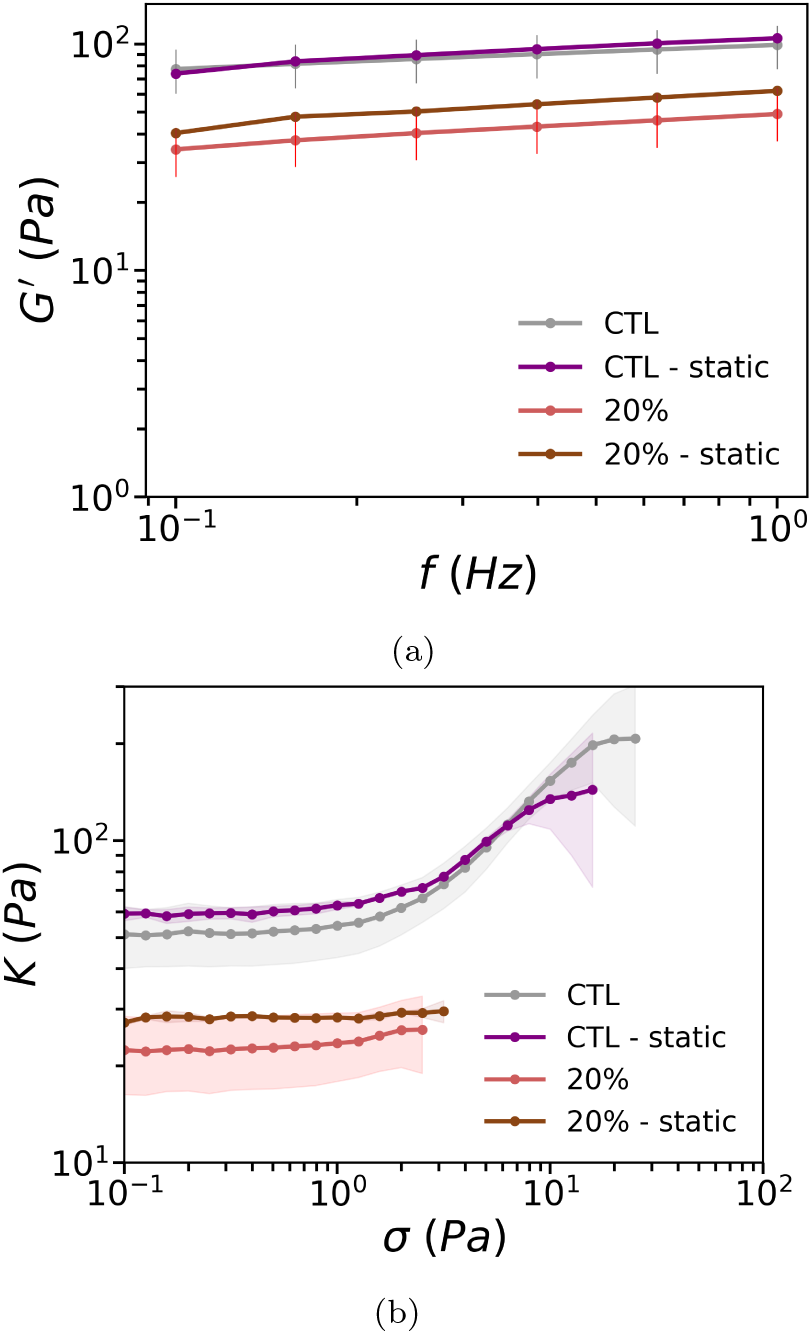
Rheological properties of collagen networks polymerized in static conditions *vs* small oscillatory strain conditions are identical. (a) Storage moduli *G*^′^ obtained from small amplitude oscillatory measurements at different frequencies for control collagen networks (CTL) and collagen networks containing cancer cells (at 20% volume fraction). (b) Corresponding differential moduli *K* as a function of applied stress stress *σ* for the final networks. Static conditions (labeled as *static*) refer to networks polymerized between the rheometer parallel plates at 37°C for 90 minutes without shearing, followed by frequency sweeps and stress ramps. The results in static conditions (CTL and 20%: n=2, N=1) are comparable to those obtained under oscillatory shear conditions (CTL: n=16, N=8 and 20%: n=16, N=8), confirming that the application of small shear oscillations during collagen polymerization does not affect the rheological properties of the network, neither for cell-free nor for cell-embedded networks. Graphs in (a) and (b) represent mean ± SD.

**Figure S7:**
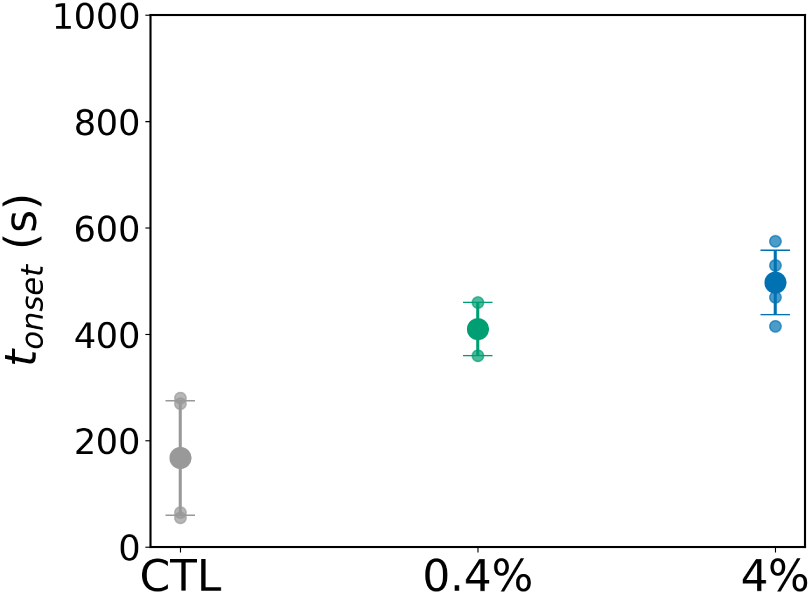
Onset of polymerization time *t_onset_* determined from the increase of the storage modulus with time during polymerization of HDF-embedded collagen networks. Cell-embedded networks containing HDF cells at volume fractions of 0.4% or 4% show a longer onset time than control networks (CTL). Data represent mean ± SD.

**Figure S8:**
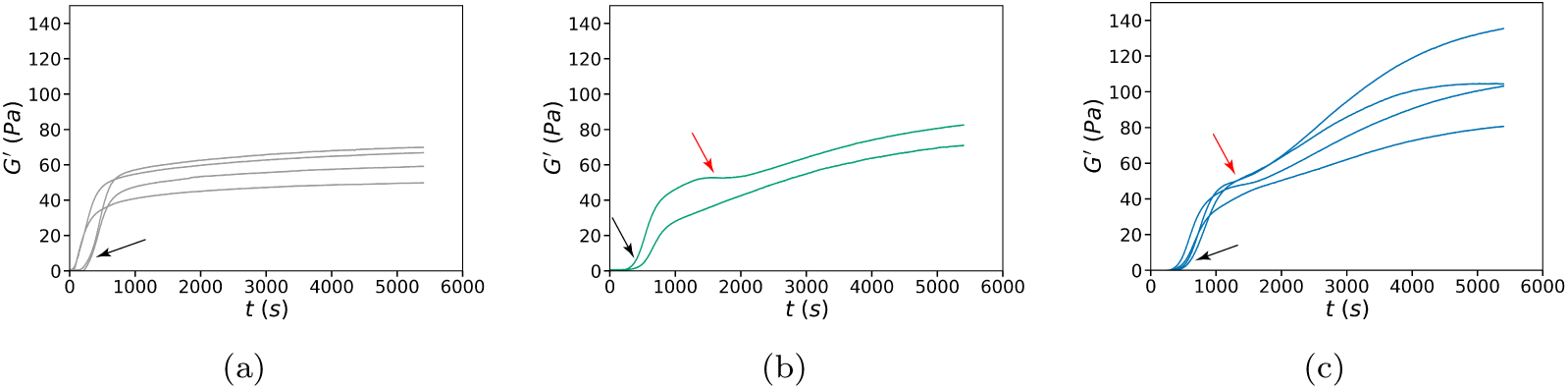
Time-dependent rheology data showing individual polymerization curves of collagen networks containing fibroblasts (HDF cells). (a) Storage modulus *G*^′^ as a function of time for control collagen (n=4, N=1). Curves show a monophasic increase of the modulus that sets in around 200 s (marked by black arrow). (b) Storage modulus *G*^′^ as a function of time for collagen containing HDF cells at a volume fraction of 0.4% (n=2, N=1). (c) Storage modulus *G*^′^ as a function of time for collagen containing HDF cells at a volume fraction of 4% (n=4, N=1). For consistency, the control collagen in (a) is prepared with the medium used for HDF cell culture. In the presence of cells, the polymerization curves tend to show a biphasic increase of the modulus, with a first onset corresponding to collagen polymerization (black arrows in b,c) and a second onset likely corresponding to force generation by the cells on the formed network (red arrows in b,c).

**Figure S9:**
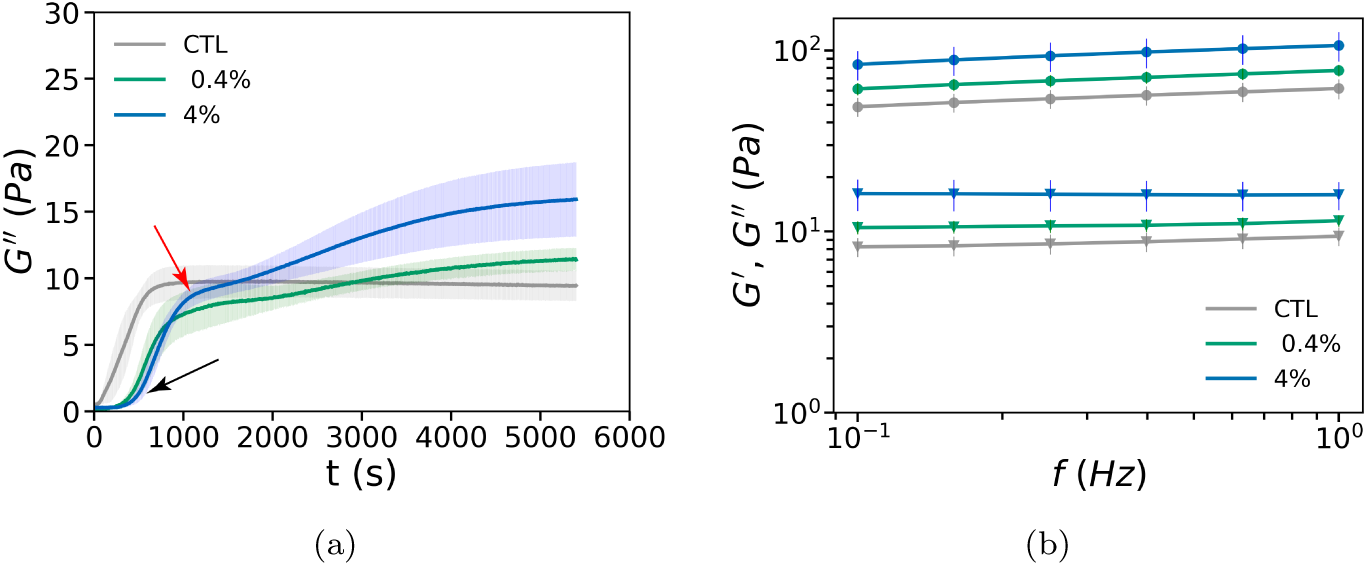
Loss modulus *G*^′^ and frequency sweeps for collagen networks containing human dermal fibroblasts (HDF). (a) Loss moduli during collagen polymerization for control collagen networks (CTL) and networks containing HDF cells at a volume fraction of 0.4% or 4%. Note that *G*^′′^ increases with time in a monophasic way for the control network, whereas the increase is biphasic in the presence of cells (black arrow and red arrow). (b) Frequency dependence of the storage moduli (circles) and loss moduli (triangles) for mature networks. Graphs represent mean ± SD.

**Figure S10:**
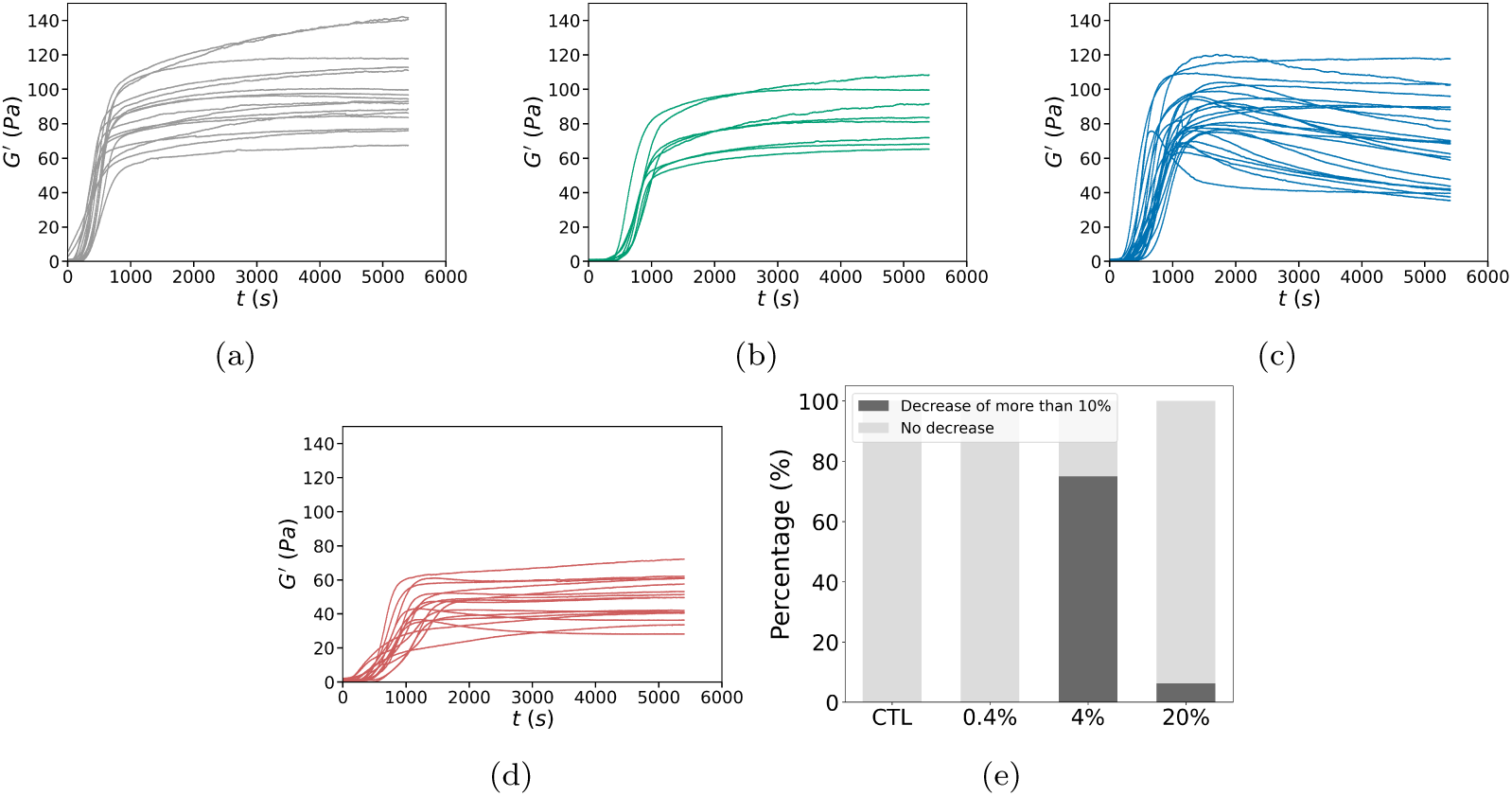
Individual polymerization curves of control and MDA-MB-231-embedded collagen matrices. Curves show the storage modulus *G*^′^ as a function of polymerization time. (a) Collagen only (n=16, N=8), prepared with MDA-MB-231 complete medium. (b-d) Cell-embedded collagen with cell volume fractions of (b) 0.4% (n=8, N=3), (c) 4% (n=24, N=10) and (d) 20% (n=16, N=8) respectively. Notably, in more than 75% of the 4% cell-embedded samples, the final *G*^′^ value, at the end of the polymerization, decreased by at least 10% from its peak value, as shown in (e).

**Figure S11:**
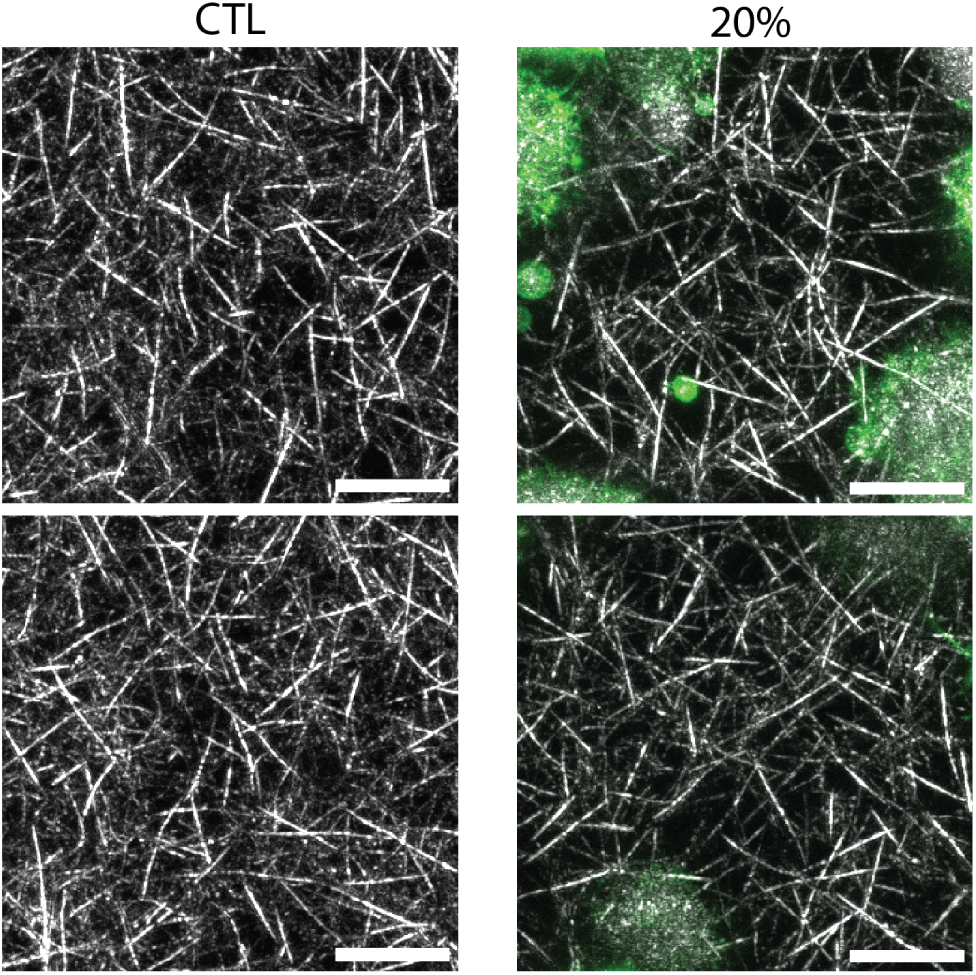
The collagen network structure is similar with and without the presence of MDA-MB-231 cancer cells. Representative maximum intensity projection images (based on confocal stacks across a depth of 10 *µ*m) of cell-free collagen hydrogels (CTL, left column) or collagen hydrogels with 20% volume fraction of MDA-MB-231 cells (20%, right column). Bottom and top rows show two different example regions of interest. Collagen fibers (grey) are imaged with reflection microscopy, while LifeAct-GFP tagged actin (green) is imaged with fluorescence microscopy. Scale bars are 10 *µ*m.

**Figure S12:**
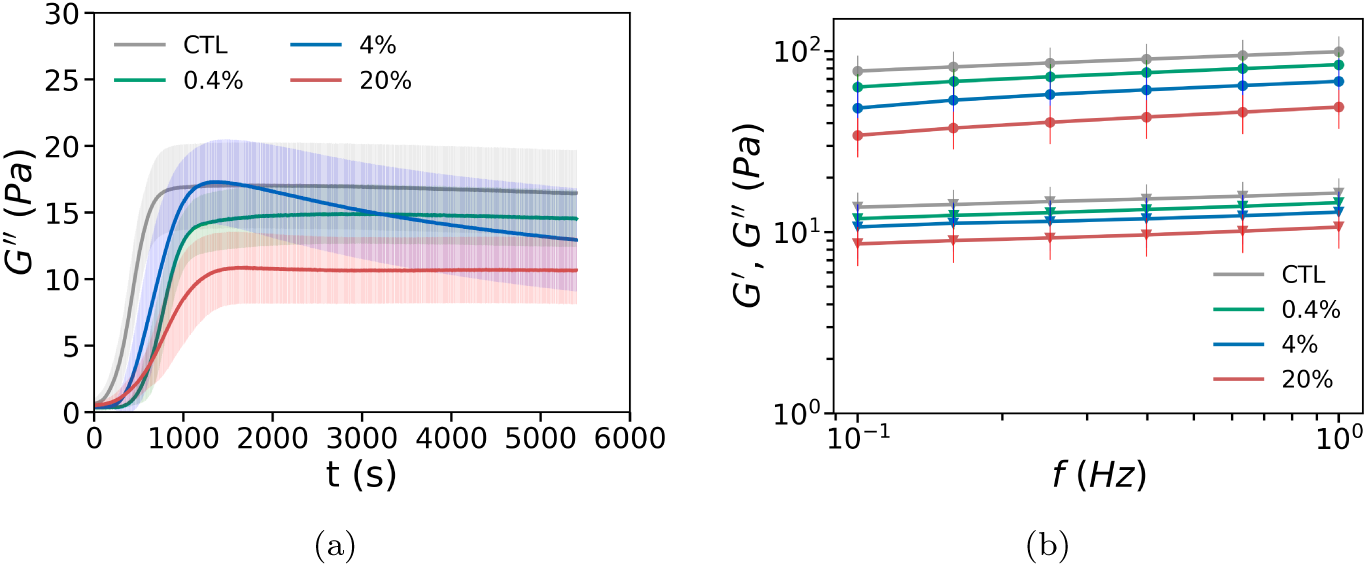
Loss modulus *G*^′′^ and frequency sweeps for collagen networks with embedded MDA-MB-231 cancer cells. (a) Loss moduli *G*^′′^ during the polymerization of collagen and MDA-MB-231-embedded collagen matrices at volume fractions of 0.4%, 4%, and 20%, showing trends similar to the storage moduli in Figure 1e of the main text. (b) Storage moduli *G*^′^ (circles) and loss moduli *G*^′′^ (triangles) from frequency sweep experiments on mature networks. Data represent mean ± SD.

**Figure S13:**
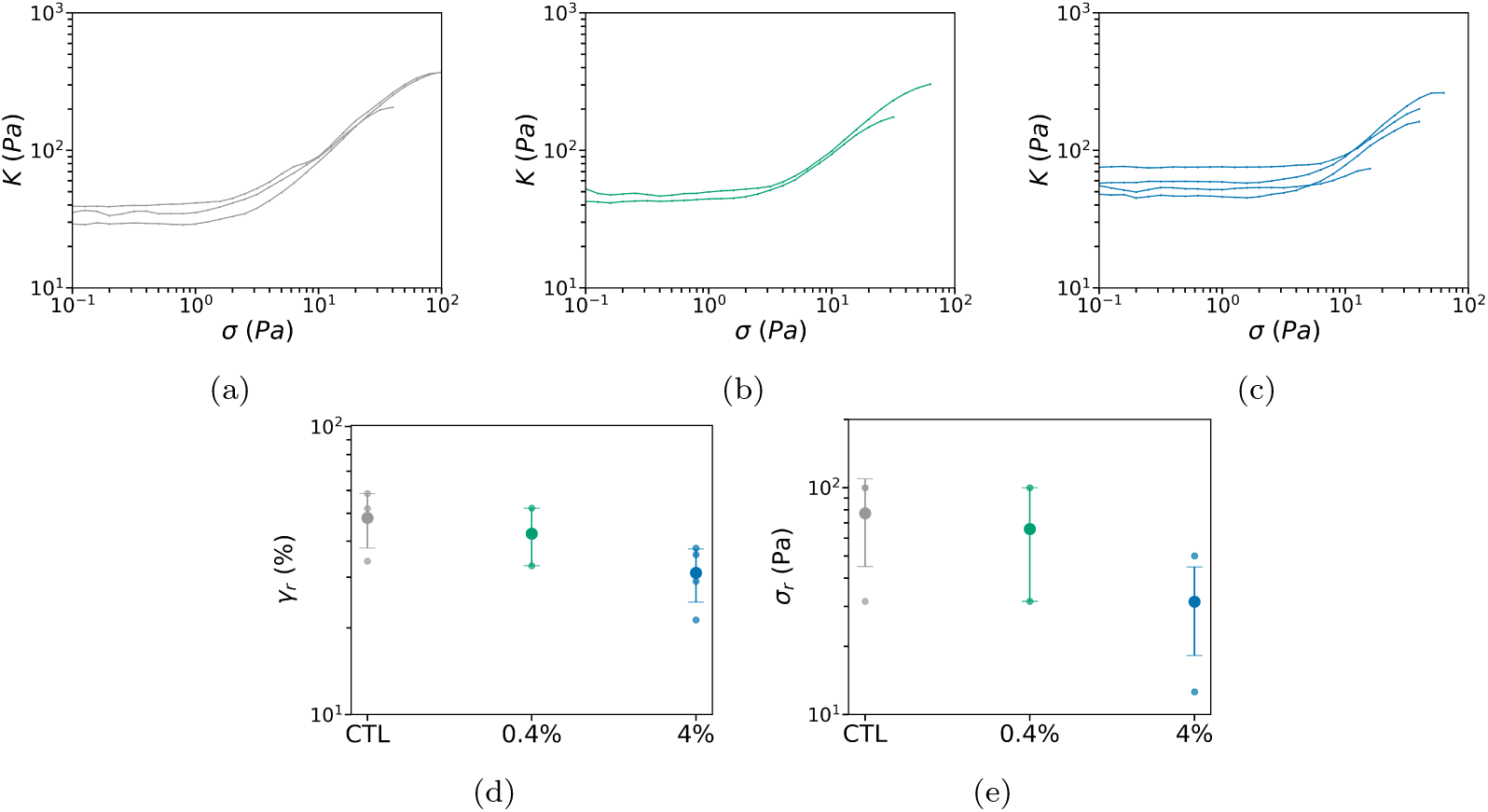
Individual stress ramp data for control collagen networks and HDF-embedded collagen matrices. (a-c) Differential modulus *K* as a function of applied shear stress *σ* for (a) collagen alone (n=3, N=1), (b) collagen with HDFs at a volume fraction of 0.4% (n=2, N=1), and (c) HDFs at a volume fraction of 4% (n=4, N=1). The curves are truncated at the rupture point. All the samples exhibit stress-stiffening behavior. (d) Strain at rupture *γ_r_*. (e) Stress at rupture *σ_r_*. Data in (d) and (e) represent mean ± SD. The strain and stress at rupture values are not significantly different between collagen control networks and cell-embedded collagen networks.

**Figure S14:**
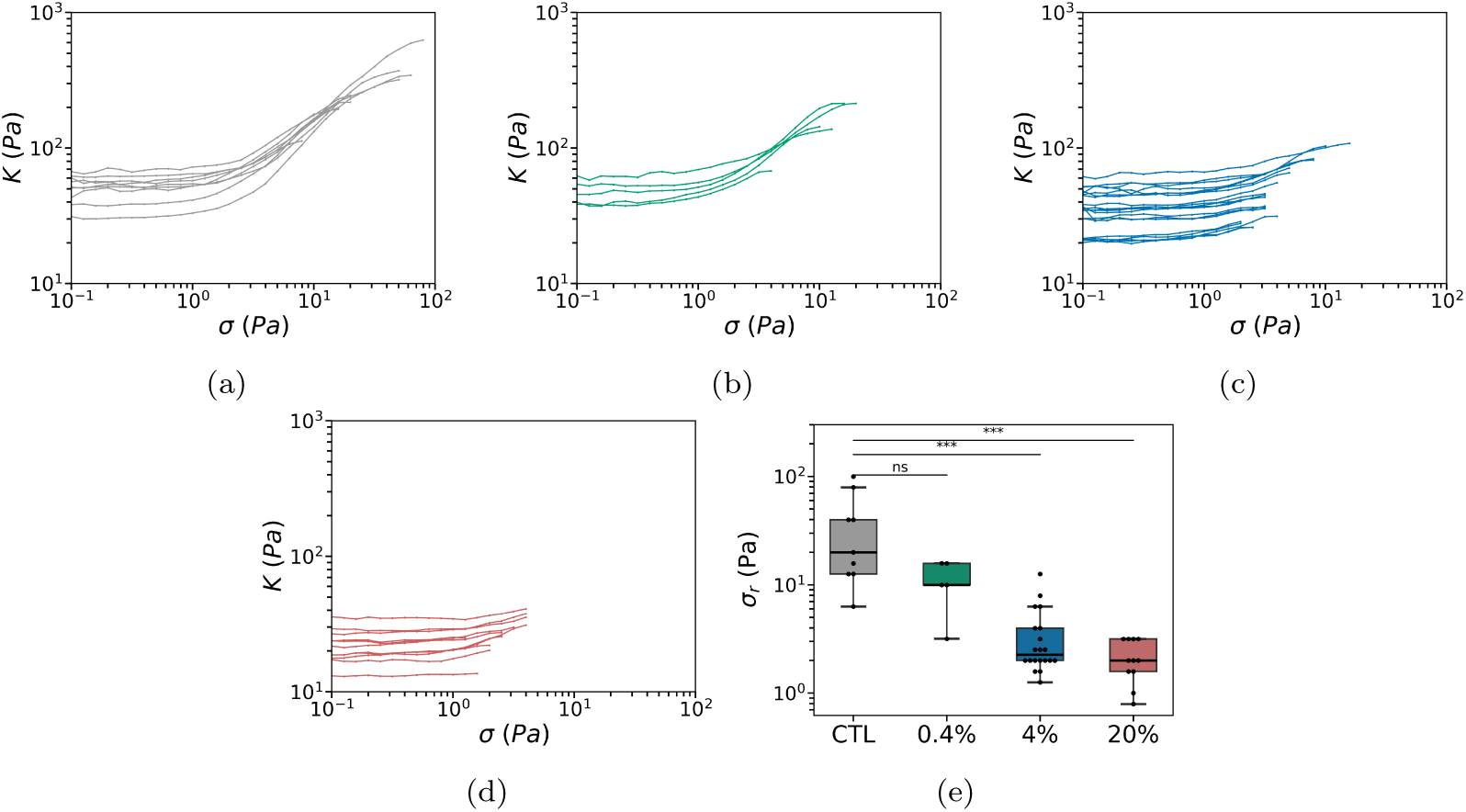
Individual stress ramp data for control collagen and MDA-MB-231-embedded collagen matrices. (a-c) Differential modulus *K* as a function of applied shear stress *σ* for (a) collagen alone (n=9, N=4), (b) collagen with MDA-MB-231 cells at a volume fraction of 0.4% (n=5, N=2), (c) 4% (n=20, N=8), and (d) 20% (n=11, N=6). The curves are truncated at the rupture point. Samples in (a) and (b) exhibit stress-stiffening behavior, while samples in (c) and (d) do not. (e) Stress at rupture *σ_r_*. Networks with 0.4% cell-embedded collagen samples behave the same as control networks, but networks with higher cell densities (4% and 20%) exhibit significantly lower values for the rupture stress compared to the control.

**Figure S15:**
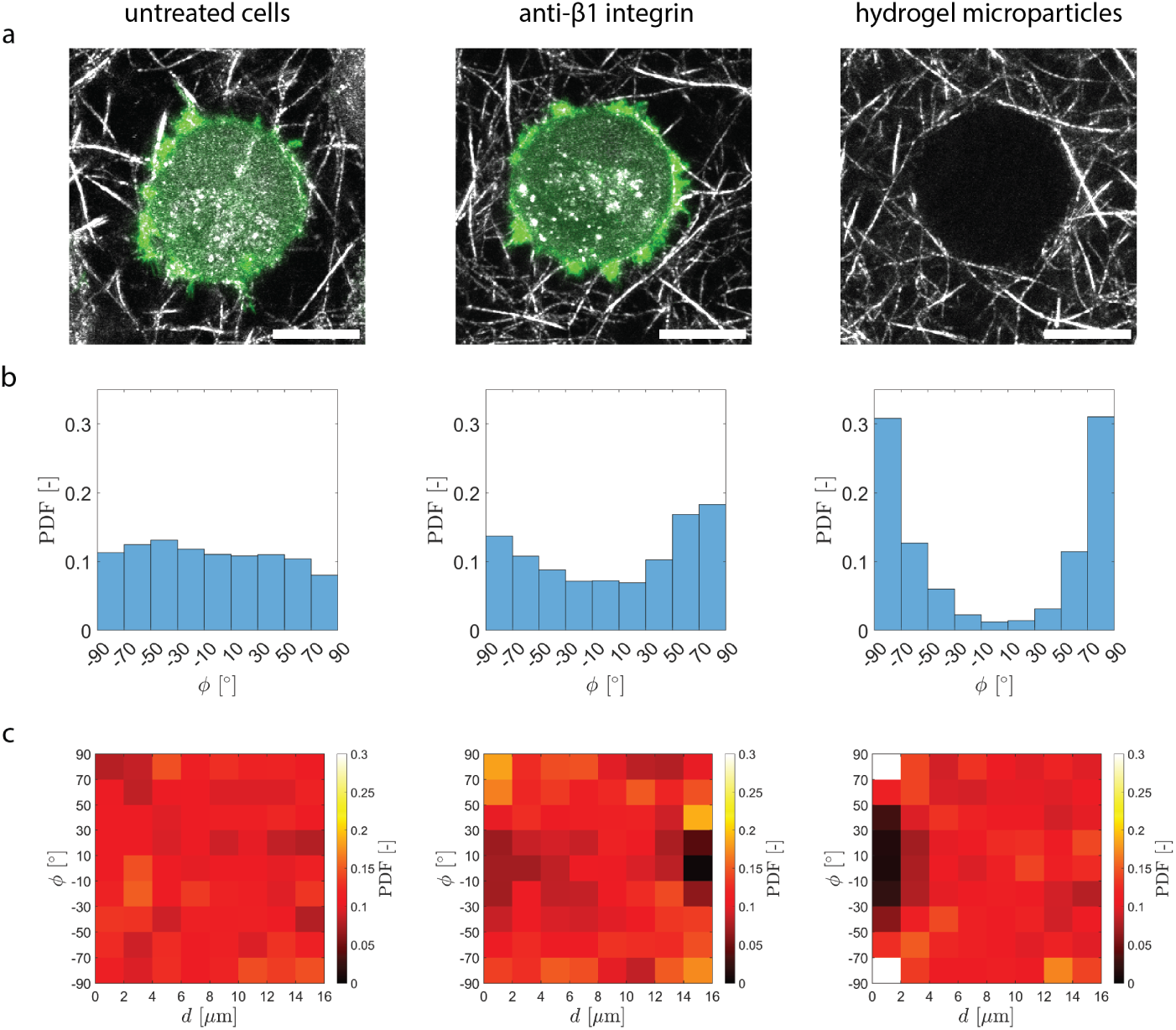
Orientation of collagen fibers around MDA-MB-231 cells or microparticles. (a) Representative maximum intensity projections (2 *µ*m) of confocal Z-stacks showing untreated cells, anti-*β*1 integrin-treated cells and hydrogel microparticles, and the surrounding collagen fibers. Collagen fibers (grey) are imaged by reflection microscopy while actin (green) is imaged by fluorescence microscopy. These images illustrate differences in fiber organization around the cells or microparticles. Scale bars are 10 *µ*m. (b) Probability density function (PDF) of the orientation (*ϕ*) of collagen fibers relative to the cell or microparticle edge and located at a distance of 2 *µ*m from the edge of untreated cells (left, n= 9, N=5), anti-*β*1 integrin-treated cells (middle, n=8, N=1), and hydrogel microparticles (right, n=10, N=1). For fibers tangential to the egde, *ϕ* = ± 90°, while for fibers perpendicular to the edge, *ϕ* = 0°. Around the hydrogel microparticles, fibers are predominantly tangentially oriented (*ϕ* = ± 90°). Anti-*β*1 integrin-treated cells exhibit more tangentially oriented fibers compared to untreated cells, which instead display an isotropic fiber distribution. (c) Heatmaps displaying the probability density function (PDF) of the orientation (*ϕ*) of collagen fibers relative to the cell or microparticle edge as a function of the distance *d* from the edge of untreated cells (left), anti-*β*1 integrin-treated cells (middle), and hydrogel microparticles (right). In the immediate proximity of the cells or microparticles, fiber orientation varies depending on the condition, as highlighted in panel (b). However, above 4 *µ*m distance from the edge, the orientation of collagen fibers becomes isotropic and randomly distributed.

**Figure S16:**
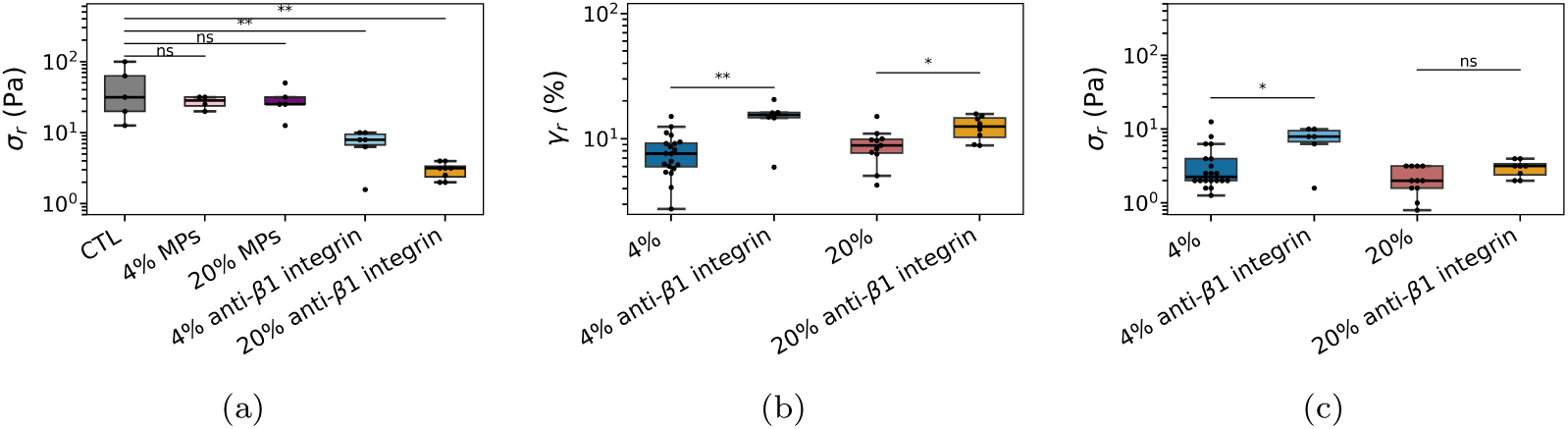
Stress at rupture for collagen alone, collagen embedded with soft hydrogel microparticles or with MDA-MB-231 cells with blocked *β*1 integrins. (a) Boxplots of the stress at rupture *σ_r_* measured for 0% (CTL), 4% and 20% volume fraction of microparticles or MDA-MB-231 cells with integrin-mediated adhesion blocked by anti-*β*1 integrin antibody. (b-c) Comparison of strain and stress at rupture of cell-embedded matrices (4% and 20% cell volume fractions) with and without blocked *β*1 integrins.

**Figure S17:**
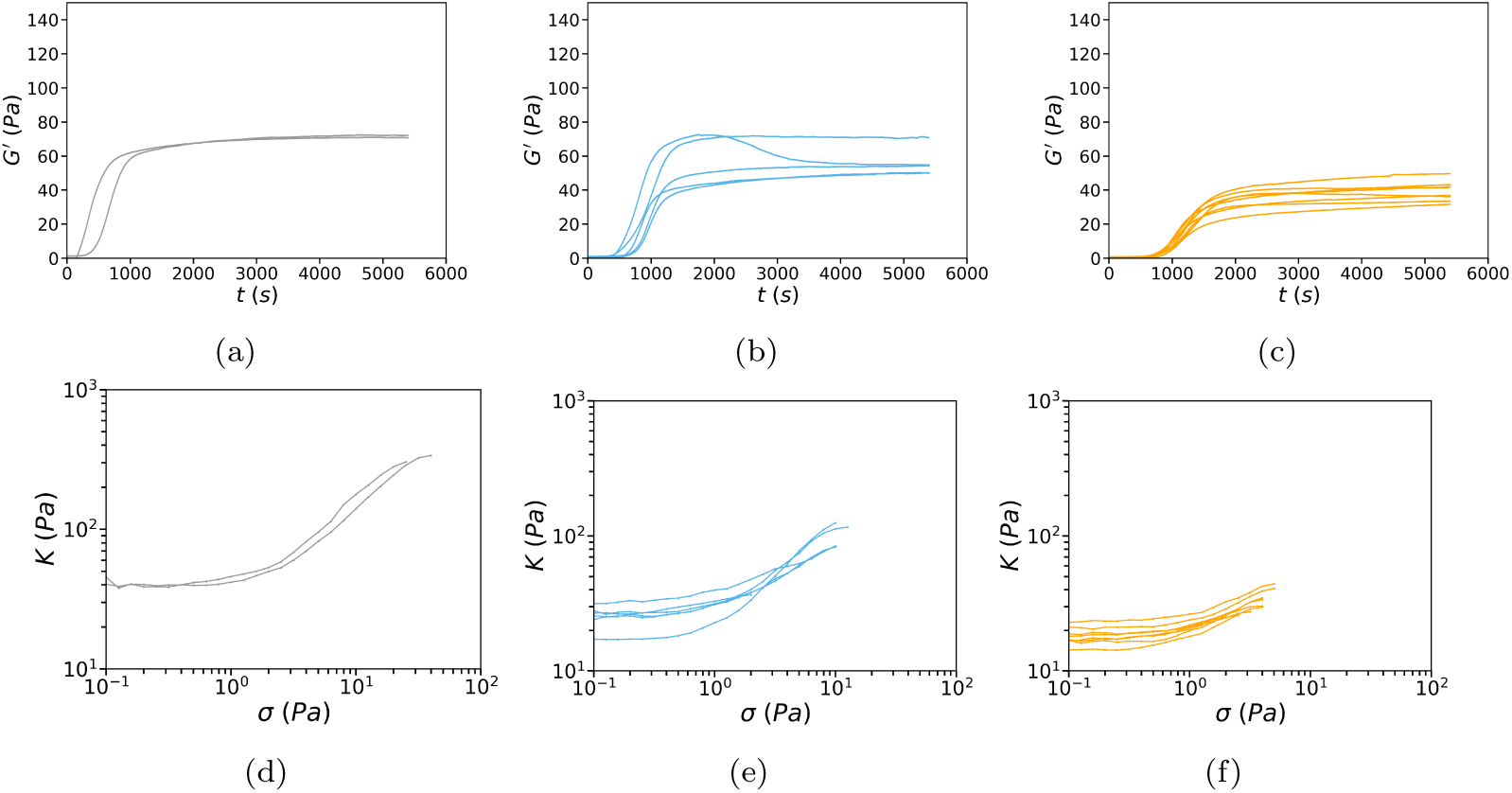
Individual polymerization curves and stress ramp data for collagen networks containing MDA-MB-231 cells with blocked *β*1 integrins. Storage modulus *G*^′^ as a function of polymerization time and differential modulus *K* as a function of the applied stress *σ* for collagen alone (n=2, N=1) (a,d), collagen with 4% cells with blocked *β*1 integrins (n=5, N=2) (b,e) and with 20% cells with blocked *β*1 integrins (n=8, N=2) (c,f). The *K* curves are truncated at the rupture point. Note that in presence of 4% cells (panel b), only one curve out of 5 shows a non-monotonic dependence of *G*^′^ on time with a peak, in contrast to samples where adhesion is not blocked (Figure S10c).

**Figure S18:**
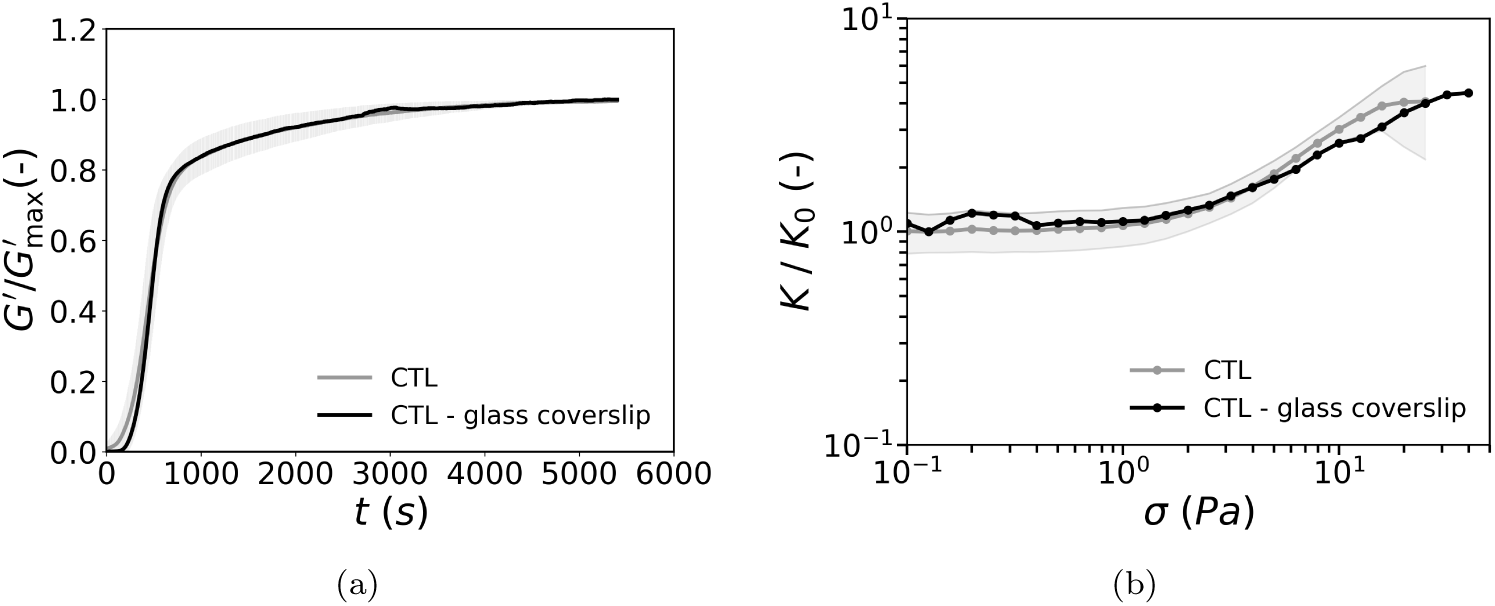
Bulk rheology of collagen networks remains unchanged when the stainless steel bottom plate is replaced by a glass coverslip. A glass coverslip (similar to the ones used for rheo-confocal microscopy) was glued on top of the bottom stainless steel plate of the rheometer. (a) Storage modulus *G*^′^ normalized by its maximal value *G*^′^_max_ as a function of polymerization time for collagen deposited on the rheometer bottom plate (grey) (n=16, N=8) and on a glass coverslip (black, n=1, N=1). (b) Differential modulus *K* normalized by the linear modulus *K*_0_ (determined as *K* at 0.1 Pa) as a function of applied shear stress *σ* for collagen deposited on the rheometer stainless steel bottom plate (grey, n=9, N=4) and on a glass coverslip (black, n=1, N=1).

**Figure S19:**
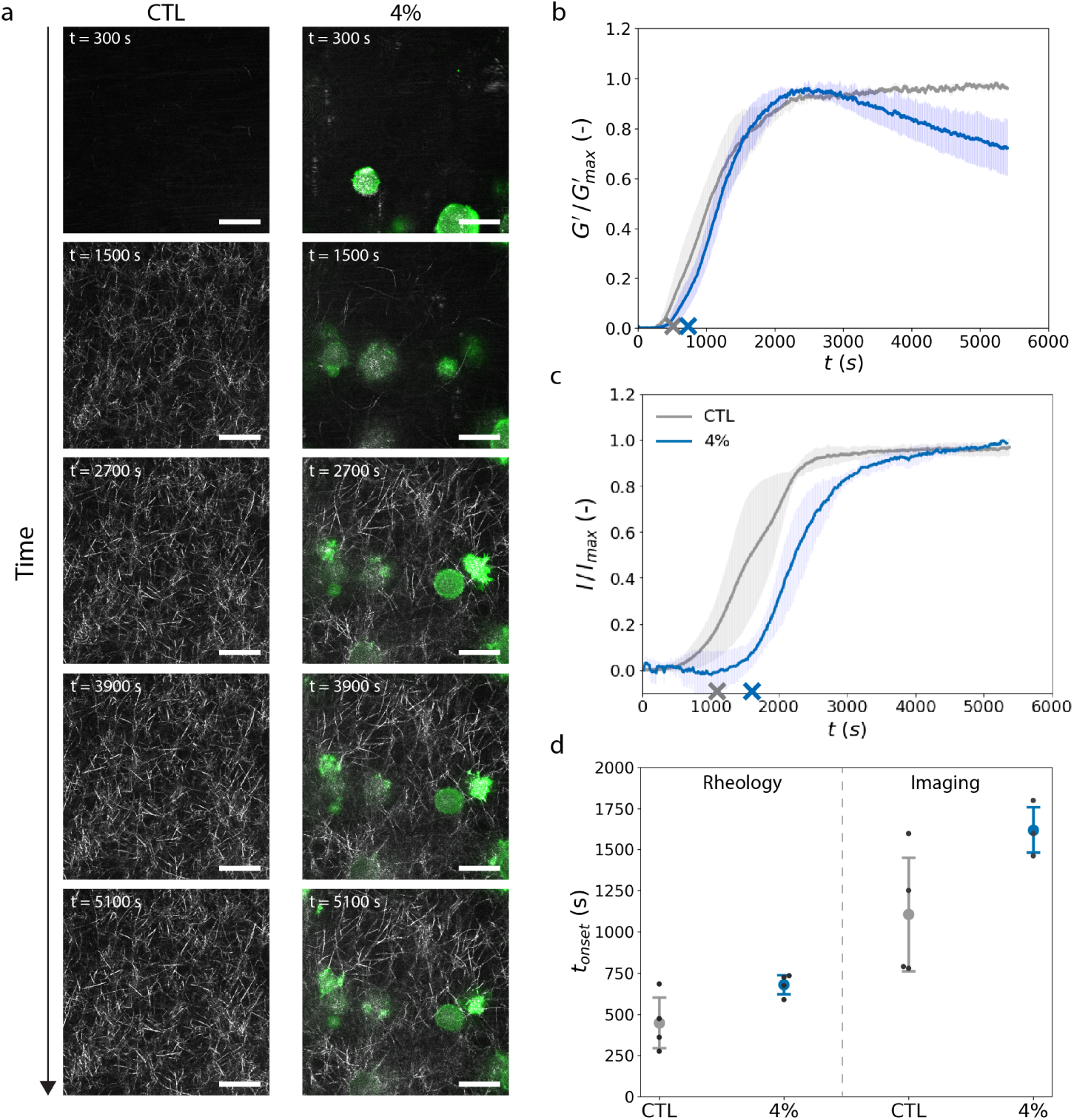
Confocal imaging and rheology measurements both demonstrate that cells delay the onset of collagen polymerization. a) Representative time lapse confocal image series acquired for collagen mixed with 0% (CTL) cells or 4% MDA-MB-231 cells during polymerization. Collagen fibers (grey) are imaged by reflection microscopy while actin (green) is imaged by fluorescence microscopy using LifeAct-GFP tagging. Scale bars are 20 *µm*. b) Storage modulus normalized by its maximal value *G*^′^ as a function of polymerization time for CTL networks and networks with 4% MDA-MB-231 embedded cells. c) Total intensity of the reflectance images of collagen normalized by the final value as a function of polymerization time for CTL networks and networks with 4% MDA-MB-231 embedded cells. Average onset times measured are indicated with crosses for CTL (grey) or 4% cells (blue) in (b) and (c). d) Onset polymerization times (*t_onset_*) for CTL or 4% MDA-MB-231 embedded cells measured from the rheological measurements in panel (b) or from the imaging data in panel (c). Data represent mean ± SD. CTL: N = 4 and 4%: N = 3.

**Figure S20:**
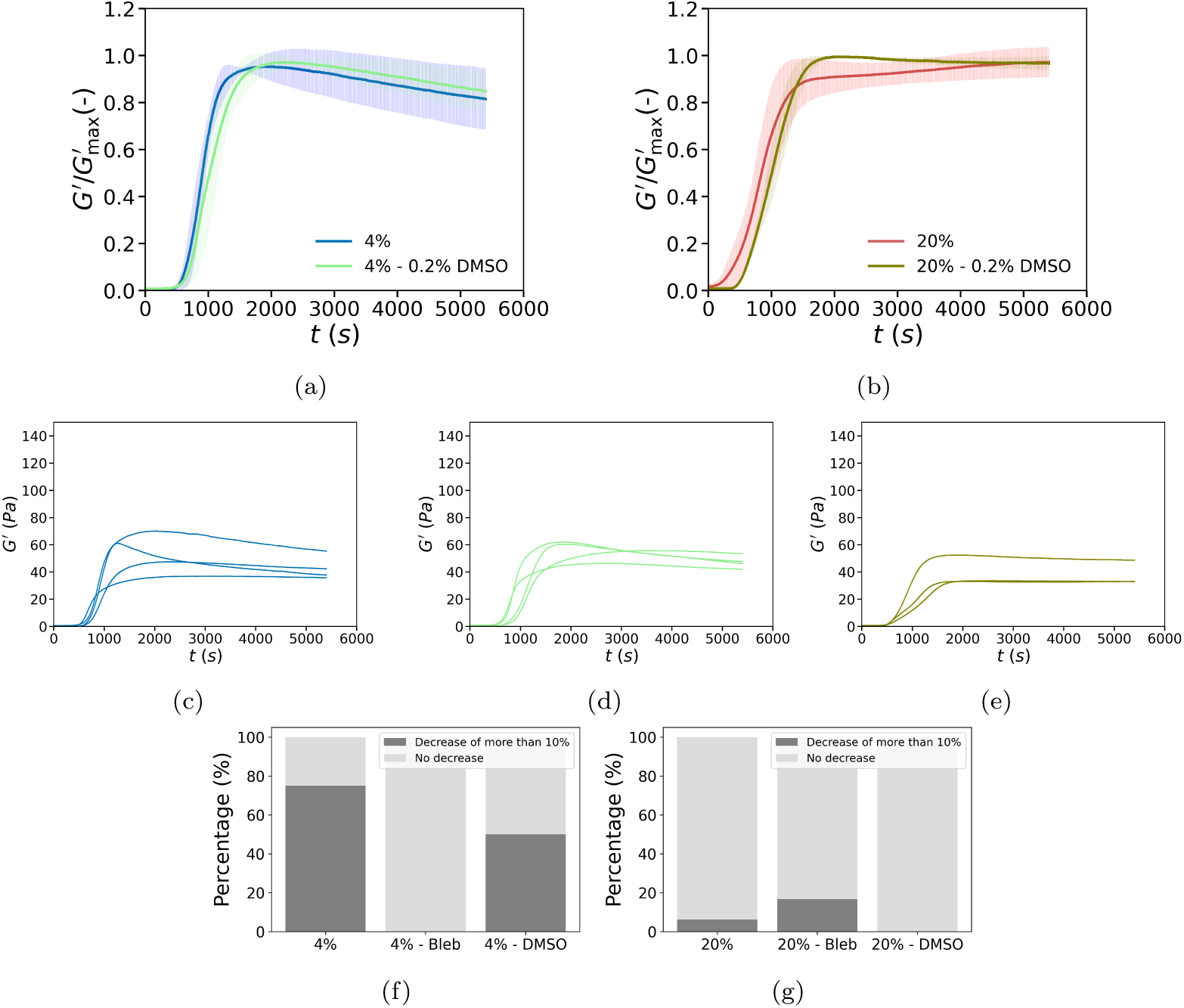
Control experiment showing that the presence of small amounts of DMSO, used as the solvent for the myosin-inhibiting drug blebbistatin, does not affect the bulk rheology of cell-embedded collagen. (a,b) Storage modulus *G*^′^ normalized by its maximum value *G*^′^ as a function of polymerization time. (a) Comparison between networks with 4% MDA-MB-231 cells exposed to 0.2% DMSO (volume percentage) for 3 hours before cell detachment and the corresponding 4% control, without DMSO. (b) Comparison between networks with 20% cells exposed to DMSO and the entire dataset for 20% cells, without DMSO. Data represent mean ± SD. (c-e) Individual curves corresponding to the averaged data from (a,b) for the control experiment with 4% cells and no DMSO (c) (n=4, N=1), 4% cells exposed to 0.2% DMSO (d) (n=4, N=1), and 20% cells exposed to 0.2% DMSO (n=3, N=2). (f,g) Histograms depicting the percentage of polymerization curves exhibiting a peak with a decrease of *G*^′^ of at least 10% compared to its peak value.

**Figure S21:**
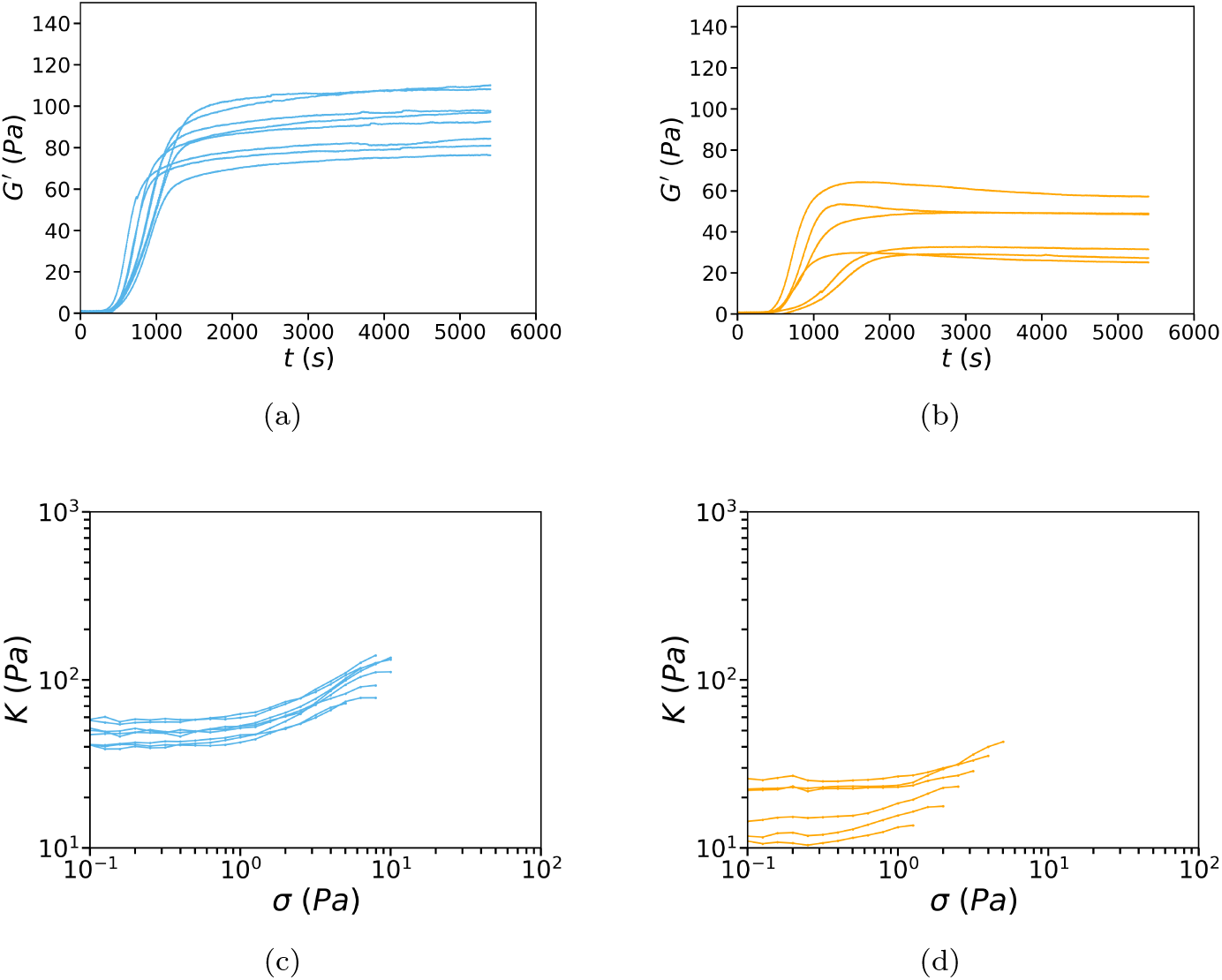
Individual polymerization curves and stress ramp data for collagen networks containing blebbistatin-treated MDA-MB-231 cells. Storage modulus *G*^′^ as a function of polymerization time and differential modulus *K* as a function of applied shear stress *σ* for collagen networks containing cells exposed to blebbistatin for 3 hours before cell passage at volume fractions of 4% (n=8, N=2) (a,c) and 20% cells (n=6, N=2) (b,d). The *K* curves are truncated at the rupture point.

**Figure S22:**
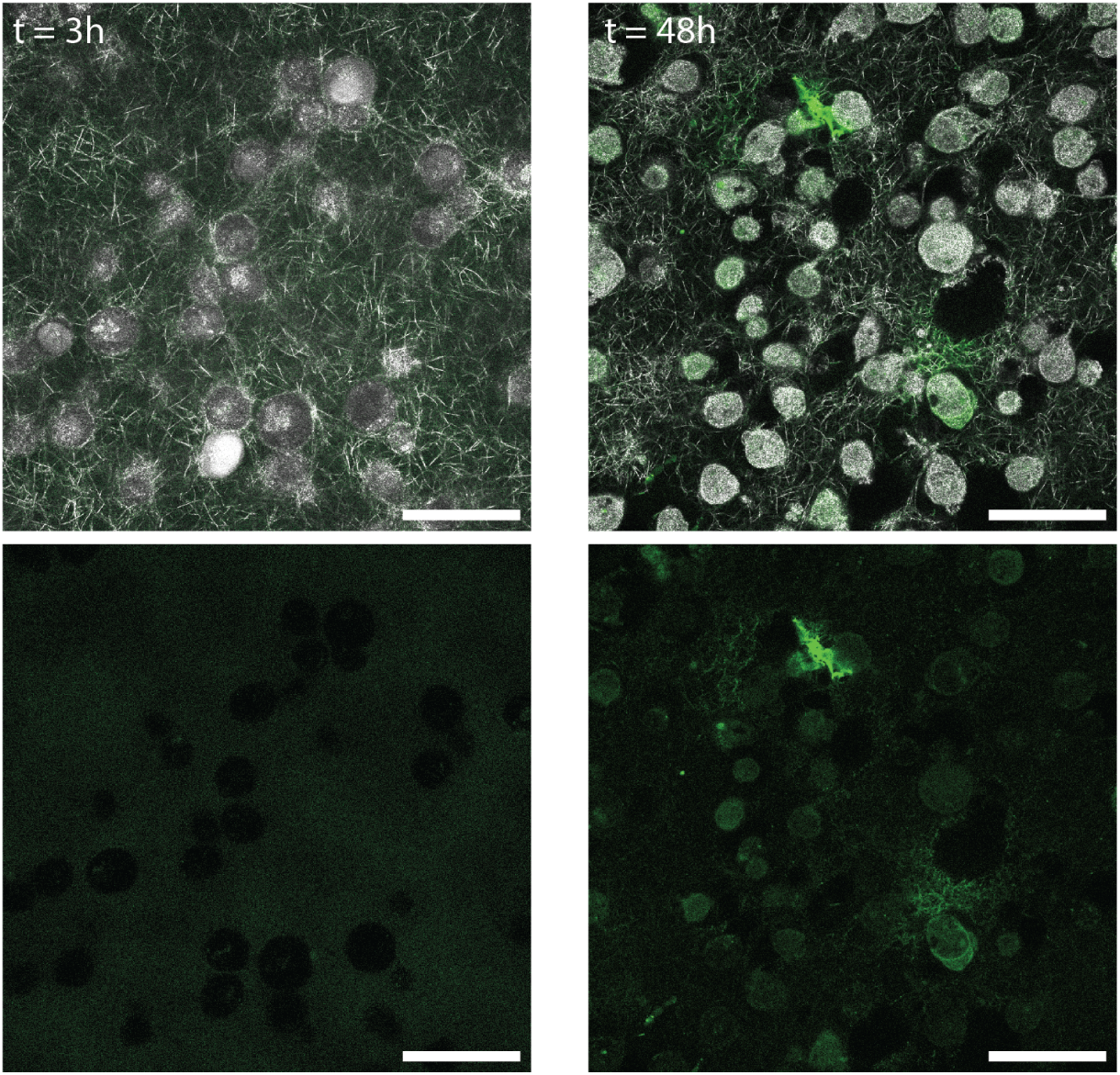
DQ-collagen assays indicate that MDA-MB-231 cells do not significantly degrade collagen on a 3 hour time scale. Collagen networks containing 25 *µ*g/mL of DQ-collagen were prepared to quantify proteolytic degradation of collagen by the cells. Collagen fibers and cells are visible in reflection microscopy (grey) while the degradation of collagen is assessed by imaging the fluorescence of DQ-collagen (green). Top panels show merged images, bottom panels show the fluorescence signal. While no degradation is observed after 3h, a clear degradation signal is visible on collagen fibers in the immediate surrounding of the cells after 48h. Scale bars are 50 *µ*m.

**Figure S23:**
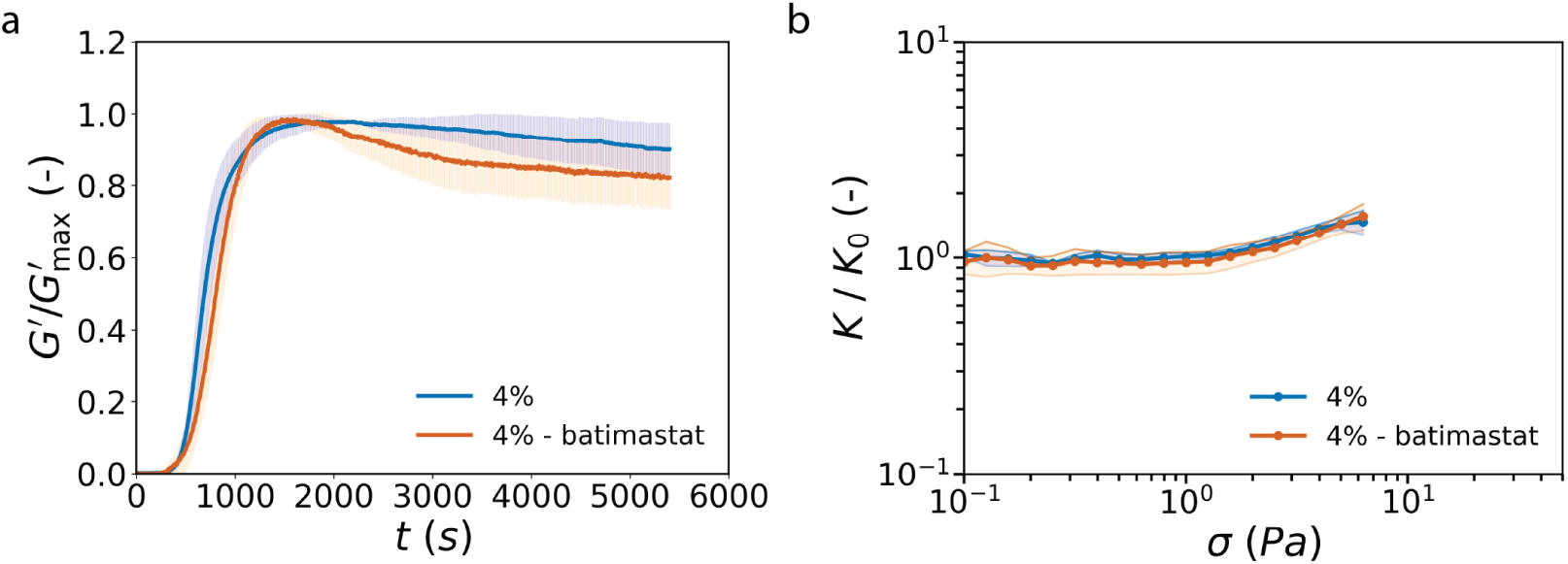
MMP inhibition with batimastat does not alter cell-mediated softening of collagen networks. a) Storage modulus *G*^′^ normalized by its maximum value G”*_max_* as a function of polymerization time. Comparison between collagen networks with 4% MDA-MB-231 cells treated with 10 *µ*M batimastat (orange; n = 3, N = 1) and untreated control samples (blue; n = 2, N = 1). A decrease of at least 10% from the peak *G*^′^ was observed in all batimastat-treated samples, comparable to the control samples. b) Differential modulus *K* normalized by the linear modulus *K*_0_ (determined as K at 0.1 Pa) for the steady-state networks with (orange, n = 2, N = 1) and without (blue, n = 2, N = 1) batimastat treatment. The non-monotonic behavior of *G*^′^ in batimastat-treated samples and the overlap of the differential moduli curves both indicate that inhibiting metalloproteinase (MMP) activity does not alter the bulk rheology of collagen networks containing cells. Thus, the observed time-dependent softening is not driven by MMP-induced matrix degradation, suggesting that other cellular processes instead underlie the softening phenomenon.

**Figure S24:**
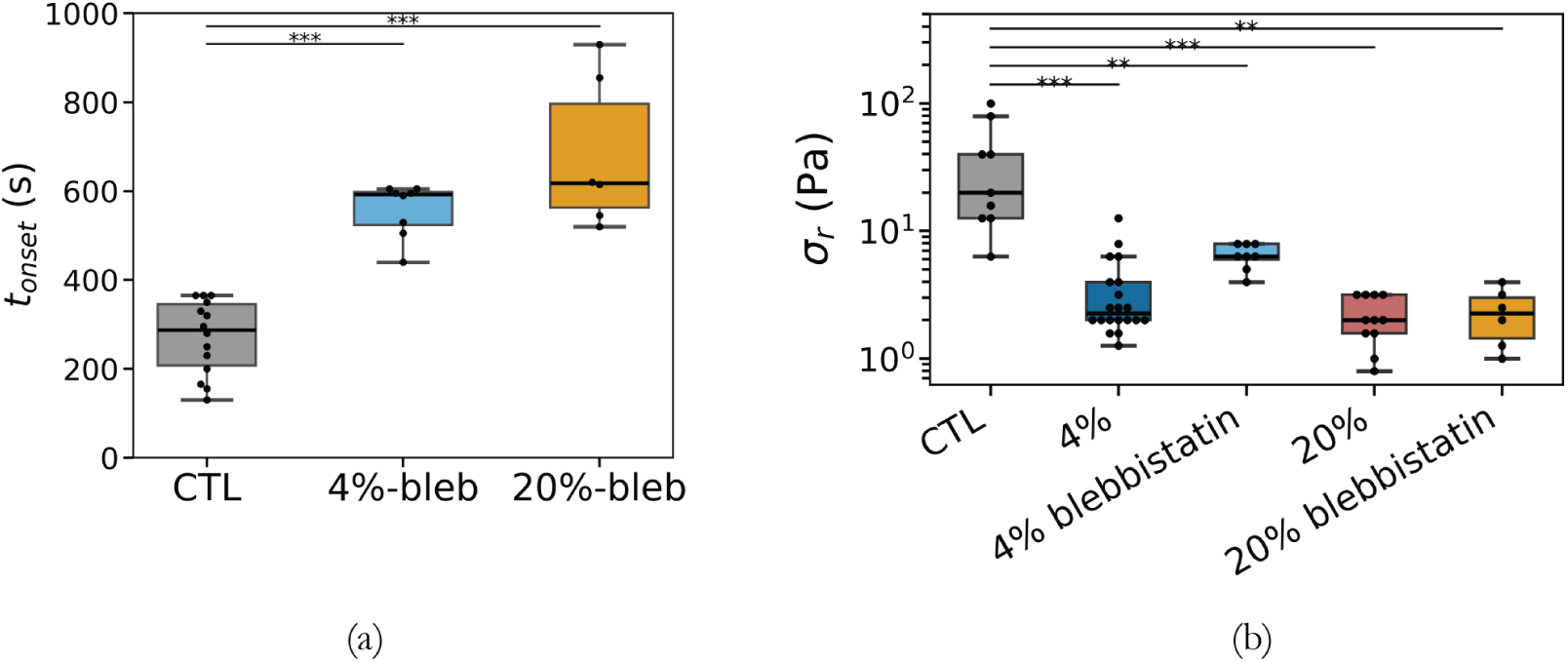
Onset time of collagen polymerization and shear stress at rupture for collagen networks containing blebbistatin-treated MDA-MB-231 cells. (a) Onset time for collagen polymerization *t_onset_* measured from rheology measurements for collagen mixed with cells exposed to blebbistatin for 3 hours before cell passage at volume fractions of 0% (CTL) (n=14, N=7), 4% (n=8, N=2), or 20% (n=6, N=2). (b) Boxplots of the stress at rupture *σ_r_*, for cell-embedded collagen with and without blebbistatin treatment.

### Supplementary Videos

**Supplementary video 1: Polymerization of a control collagen network imaged in the rheo-confocal setup.** Collagen fibers appearing over time (grey) are imaged by reflection microscopy. The visible oscillations over time are caused by the constant application of shear strain oscillations at a frequency of 1 *Hz* and amplitude of 1%.

**Supplementary video 2: Polymerization of a collagen network in presence of 4% volume fraction of MDA-MB-231 cells imaged in the rheo-confocal setup.** Collagen fibers (grey) appearing over time are imaged by reflection microscopy and actin (green) is imaged by fluorescence microscopy. The visible oscillations over time are caused by the constant application of shear strain oscillations at a frequency of 1 *Hz* and amplitude of 1%.

**Supplementary video 3: Confocal imaging of cell-matrix interactions during polymerization of collagen in the presence of 4% MDA-MB-231 cancer cells acquired using the rheo-confocal setup.** White arrows indicate cell-mediated remodeling events. Collagen fibers (grey) are imaged by reflection microscopy while actin (green) is imaged by fluorescence microscopy. The central arrow points at a cell protrusion pulling the fiber network and then retracting, while the top arrow indicates a fiber being bent by a cell.

**Supplementary video 4: Confocal imaging of cell-matrix interactions during polymerization of collagen in the presence of 4% MDA-MB-231 cancer cells acquired using the rheo-confocal setup.** White arrows indicate cell-mediated remodeling events. Collagen fibers (grey) are imaged by reflection microscopy while actin (green) is imaged by fluorescence microscopy. The arrow on the left indicates a cell protrusion that first displaces collagen fibers and then pulls on one individual collagen fiber. This pulling leads to the displacement of a crosslink node and to the bending of a fiber, indicated by the arrow on the right.

**Supplementary video 5: Confocal imaging of cell-matrix interactions during polymerization of collagen in the presence of 4% MDA-MB-231 cancer cells acquired using the rheo-confocal setup.** White arrows indicate cell-mediated remodeling events. Collagen fibers (grey) are imaged by reflection microscopy while actin (green) is imaged by fluorescence microscopy. Arrows indicate cell protrusions that displace collagen fibers and locally remodel the structure of the network.

**Supplementary video 6: Confocal imaging during the shear stress ramp of a control collagen network acquired using the rheo-confocal setup.**A stress ramp is applied to the mature collagen network with stresses logarithmically increasing from 0.01 Pa to 100 Pa at a rate of 10 points per decade and with 5 s between each point. Collagen fibers (grey) are imaged by reflection microscopy. At early times corresponding to low stresses, there is no visible movement of the sheared network. After about 100 s, corresponding to an applied stress of 1 Pa, we observe a global displacement of the network to the right, corresponding to the direction of shear. The disappearance of the reflection signal from the collagen network for the final frames corresponds to the rupture point.

**Supplementary video 7: Confocal imaging during the shear stress ramp of collagen in the presence of 4% MDA-MB-231 cancer cells acquired using the rheo-confocal setup.** A stress ramp is applied to the mature network with stresses logarithmically increasing from 0.01 Pa to 100 Pa at a rate of 10 points per decade and with 5 s between each point. Collagen fibers (grey) are imaged by reflection microscopy while actin (green) is imaged by fluorescence microscopy. A global displacement of the network in the direction of shear is visible. The collagen network and the cells seem to translate laterally in a similar manner. No clear differences in the local displacement of collagen fibers are evidenced when comparing to the stress ramp videos of control collagen network. The final frame of the video corresponds to the rupture point, indicated by the loss of actin fluorescence signal.

**Supplementary video 8: Polymerization of collagen network in the presence of 20% volume fraction of microparticles (MPs) imaged in the rheo-confocal setup.** Collagen fibers (grey) appearing over time are imaged by reflection microscopy, while the MPs appear as voids within collagen fibers. The visible oscillations over time are caused by the constant application of shear strain oscillations at a frequency of 1 Hz and amplitude of 1%. Collagen fibers appear in random locations and no preferential nucleation of collagen fibers around the microparticles can be observed.

**Supplementary video 9: Confocal imaging during the shear stress ramp in the presence of 20% volume fraction of microparticles (MPs) imaged in the rheo-confocal setup.** A stress ramp is applied to the mature network with stresses logarithmically increasing from 0.01 Pa to 100 Pa at a rate of 10 points per decade and with 5 s between each point. Collagen fibers (grey) are imaged by reflection microscopy, while the MPs appear as voids within collagen fibers. A global displacement of the network in the direction of shear is visible. The collagen network and the MPs seem to translate laterally in a similar manner, with no deformation of the MPs up to about 10 Pa (corresponding to time point 2:30 of the stress ramp). After this point and right before rupture, the holes in the collagen network seem to expand. The final frame of the video corresponds to the rupture point, corresponding to 20 Pa in this example.

